# *Muc6*-expressing gastric isthmus progenitors contribute to regeneration and metaplasia supported by myeloid-mesenchymal interactions

**DOI:** 10.1101/2025.04.15.648856

**Authors:** Junya Arai, Yusuke Iwata, Ayumu Tsubosaka, Hiroto Kinoshita, Shintaro Shinohara, Sohei Abe, Toshiro Shiokawa, Katsuyuki Oura, Nobumi Suzuki, Masahiro Hata, Ken Kurokawa, Yukiko Oya, Mayo Tsuboi, Sozaburo Ihara, Keita Murakami, Chihiro Shiomi, Chie Uekura, Hiroaki Fujiwara, Hiroaki Tateno, Seiya Mizuno, Satoru Takahashi, Yutaka Suzuki, Akinori Kanai, Tetsuo Ushiku, Hideaki Ijichi, Yoshihiro Hirata, Masato Kasuga, Valerie P. O’Brien, Nina Salama, Miwako Kakiuchi, Shumpei Ishikawa, Timothy C. Wang, Yoku Hayakawa, Mitsuhiro Fujishiro

**Author notes:** **Corresponding author:** Yoku Hayakawa, Department of Gastroenterology, Graduate School of Medicine, University of Tokyo, Hongo, Bunkyo-ku, Tokyo 113-8655, Japan. Co-1^st^ authors (Junya Arai, Yusuke Iwata).

## Abstract

Gastric mucosal homeostasis is maintained by tissue-resident stem and progenitor cells residing in the isthmus region. Following mucosal injury, surviving cells contribute to regeneration, coinciding with characteristic pathological changes such as atrophic gastritis and metaplasia. To comprehensively understand the cellular dynamics involved in this process, we performed single-cell and spatial transcriptomics using newly generated transgenic mice. In human samples and mouse models, loss of gastric chief cells precedes, and even induces, loss of parietal cells during the progression of atrophy and metaplasia, validating the causal relationship underlying the decrease of these two lineages. Single-cell analysis confirmed robust stemness and metaplastic changes in the *Muc6*-expressing neck lineage following either chief or parietal cell ablation, and lineage-tracing experiments revealed that *Muc6*-expressing isthmus progenitors serve as a source of metaplasia and regeneration. Mechanistically, mucosal injury recruits IL-1-expressing myeloid cells, which stimulates NRG1 production in stromal fibroblasts, leading to mucosal proliferation and regeneration mediated by Myc activation in isthmus progenitors. These findings highlight the injury-responsible stem cell-like function of *Muc6*-expressing isthmal progenitors, which play a critical role in mucosal homeostasis and disease progression.

**Visual abstract:** 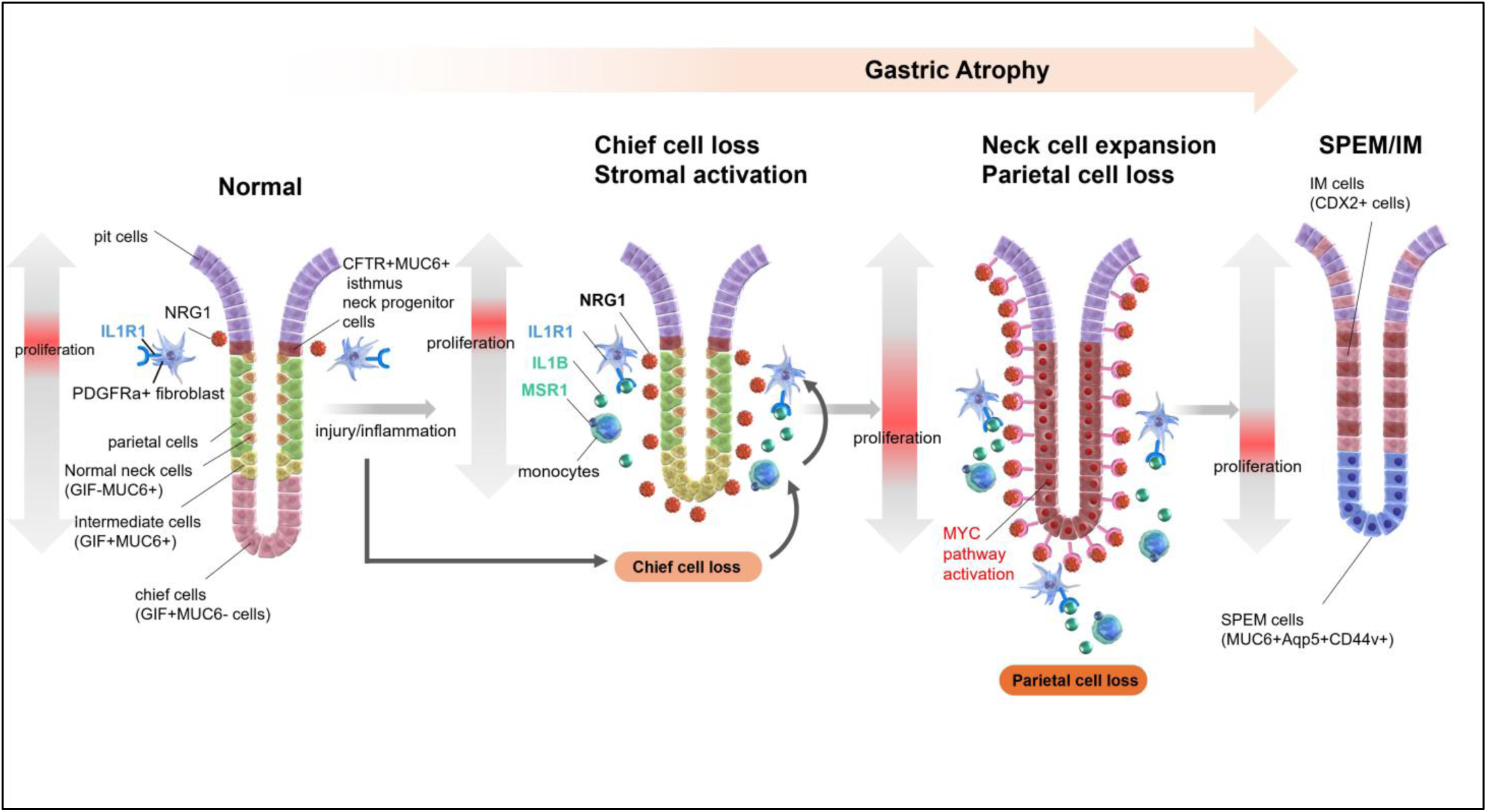

## Introduction

Gastric cancer (GC) remains a leading cause of cancer-related mortality worldwide^1^. Chronic infection with *Helicobacter. pylori*, the primary gastric carcinogen, causes characteristic pathological changes in the stomach, namely gastric atrophy and metaplasia, which drive the multistep carcinogenic cascade known as the Correa pathway^2^. It is hypothesized that chronic inflammation and mucosal injury disrupt normal cell cycle and turnover during tissue repair, leading to the accumulation of genetic and epigenetic changes in the long-lived stem cells. These changes in stem cells likely contribute to precancerous pathological transformations. However, the precise mechanisms and cellular behaviors underlying these stepwise alterations in the stomach remain poorly understood.

The gastric epithelium undergoes continuous turnover, maintained by long-lived stem cells that generate daughter cell lineages^3^. The oxyntic stomach, also known as the corpus, constitutes the main body of the stomach and harbors a variety of mature cell types essential for digestion and mucosal protection (Figure S1A). These include acid-secreting parietal cells, mucous-producing surface pit and mid-glandular neck cells, and pepsinogen (PG)-secreting gastric chief cells. In the normal corpus, stem cells and their immediate progenitors reside in the isthmus region, located in the upper third of the glands. Several stem cell markers such as *Stmn1* or *Mist1* have been reported in the corpus^4,5^, while specific markers for gastric progenitor cells, besides broad proliferation markers such as *Mki67* or *Pcna*, have not been identified. They generate MUC5AC+ pit cells that migrate towards the mucosal surface, while simultaneously producing H/K-ATPase+ parietal cells and MUC6+ neck cells that migrate towards the gland base. During gland turnover and cell migration, neck cells eventually differentiate into PG+ chief cells^6^. However, gastric injury and inflammation disrupt this normal differentiation process, leading to a marked reduction in two secretory lineages: parietal and chief cells, characteristic of the atrophic stomach. Furthermore, aberrant cell lineages—normally absent in the stomach—frequently emerge following gastric atrophy, a phenomenon collectively referred to as gastric metaplasia^7^. The presence and degree of atrophy and metaplasia have been shown to be closely associated with gastric cancer incidence in human cohorts^8,9^.

Previous studies, primarily conducted in murine models, have investigated the origins and mechanisms of regeneration and metaplasia following acute gastric mucosal injury. Most researchers have utilized pharmacological injury models, which rapidly induce parietal cell loss and the development of spasmolytic polypeptide-expressing metaplasia (SPEM)^10^. SPEM is a form of gastric metaplasia characterized by the abundant expression of neck cell markers, such as TFF2 and MUC6, as well as the ectopic expression of gastric antral gland markers, including CD44 and AQP5. Given that a subset of these metaplastic cells expresses low levels of chief cell markers, some studies suggest that gastric metaplasia originates from gastric chief cells following parietal cell loss^11–16^. However, it is important to note that reagents used to induce gastric injury also significantly impair and reduce the chief cell population. Additionally, there is a marked expansion of the proliferative isthmus zone following injury, supporting the prevailing hypothesis that metaplasia and gland regeneration primarily originate from isthmus stem cells^3,17–22^. Nevertheless, the causal relationship between impaired cell differentiation and metaplasia, as well as the ultimate origin of metaplastic cells, remains a topic of ongoing debate^17,19,23–25^.

To address these gaps in knowledge, we established genetically engineered mouse models designed to induce the selective ablation of chief and parietal cells. We conducted comprehensive transcriptomic analyses and *in vivo* lineage-tracing studies. By integrating data from human tissue samples, we elucidated the molecular mechanisms and cellular dynamics underlying gastric injury and repair, particularly in relation to metaplasia development.

## Result

### Chief cell loss precedes and induces parietal cell loss, leading to the expansion of metaplasia

To investigate the sequential changes in chief cells, parietal cells, and metaplastic cells during gastric atrophy, we analyzed 72 human stomach samples at various stages of atrophy. Patients were selected based on matched severity levels of atrophic gastritis, age, sex, and history of gastric cancer (GC), as outlined in the flowchart (Figure S1B). Immunohistochemical (IHC) staining revealed a progressive reduction in H/K-ATPase+ parietal cells and PG-I+ chief cells, along with an increase in Ki67+ proliferative cells and TFF2+CD44v9+ metaplastic cells, correlating with the severity of gastric atrophy (Figure 1A-B). Notably, a decline in chief cells was observed earlier than the reduction in parietal cells, coinciding with increased cellular proliferation (Figure 1C and S1C). Similar trends were observed when patients were stratified based on intestinal metaplasia (IM) grades (Figure S1D-E). We then infected mice with *H. pylori* and investigated histological changes along the time course. Consistent with human data, a decrease in chief cells was observed earlier than that in parietal cells (Figure S1F). These findings indicate that in humans and mice, chief cell loss precedes parietal cell loss during *H. pylori*-induced gastric atrophy and metaplasia development.

**Figure 1.**
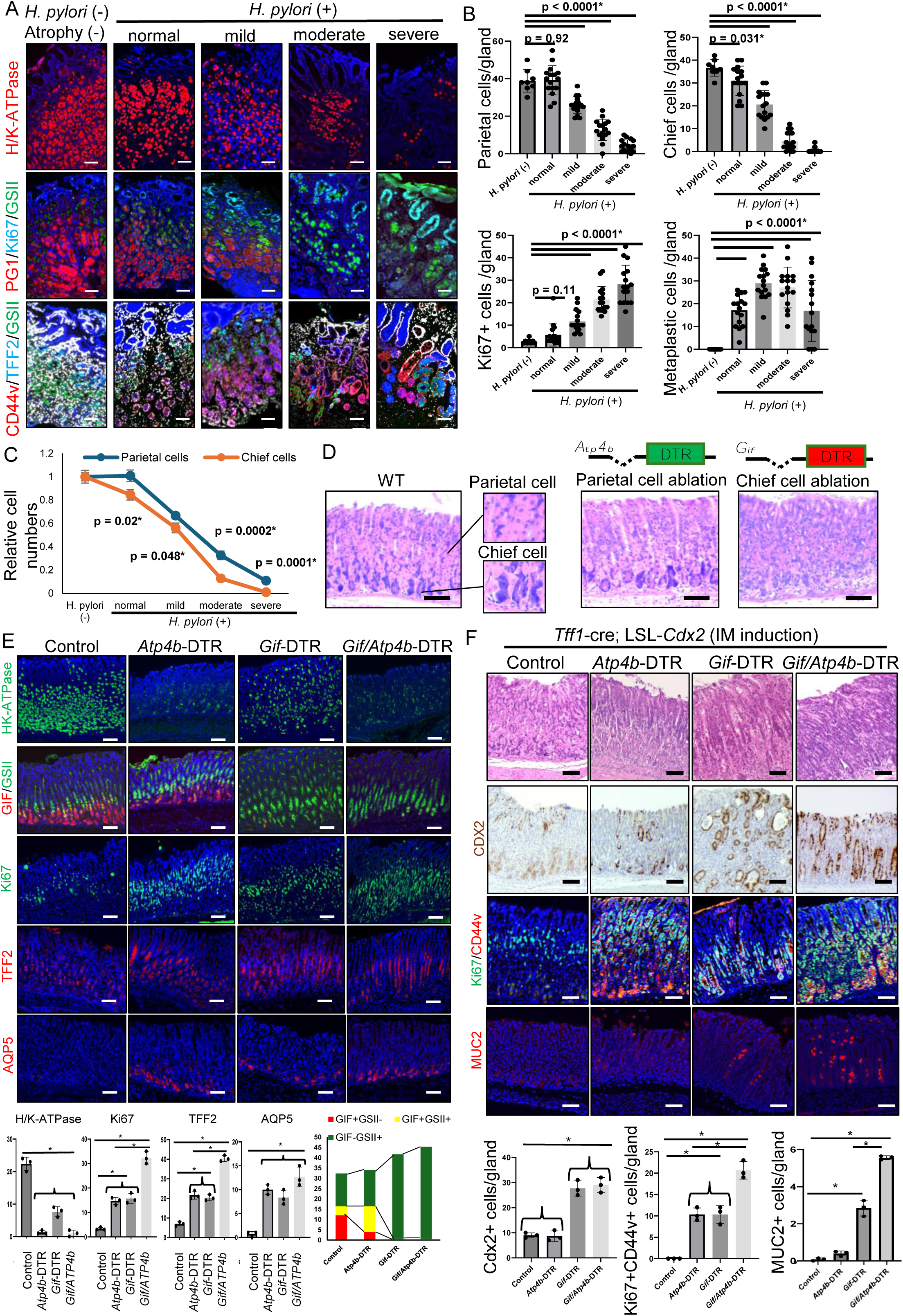
Chief cell loss precedes and induces parietal cell loss, leading to the expansion of metaplasia. (A-C) H/K-ATPase, pepsinogen (red)/Ki67 (light blue)/GSII (green), and CD44v (red)/TFF2 (light blue)/GSII (green) staining in human samples selected according to the severity of gastric atrophy. (A) Representative images. (B) H/K-ATPase+ parietal cells, pepsinogen I+ chief cells, Ki67+ proliferative cells, and TFF2+/CD44v+ metaplastic cells per gland were quantified. (C) The comparison of the number of chief cells and Ki67 + cells shown as scatter diagram. (D) (left) HE images of WT mouse stomach (eosinophilic parietal cells and basophilic chief cells are enlarged). (middle and right) Gene constructs of *Atp4b*-DTR and *Gif*-DTR mice, and HE images of these mice one day after DT treatment. (E) H/K-ATPase (green), GIF (red)/GSII (green), Ki67 (green), TFF2 (red), and AQP5 (red) staining and numbers of positive cells per gland in WT, *Atp4b*-DTR, *Gif*-DTR, *Gif*-DTR; *Atp4b*-DTR mice three days after DT treatment (n=3 mice/group). (F) Cdx2 (brown), Ki67 (green)/CD44v (red), and MUC2 (red) staining and numbers of positive cells per gland in *Tff1*-cre; LSL-*Cdx2, Tff1*-cre; LSL-*Cdx2*;*Gif*-DTR, *Tff1*-cre; LSL-*Cdx2*; *Atp4b*-DTR, *Tff1*-cre; LSL-*Cdx2*; *Gif*-DTR; *Atp4b*-DTR mice (n=3 mice/group). Mice were treated with DT three times per week for two weeks. Scale bars; 100 μm. Mean ± S.E.M. *P < .05.

To determine the causal relationship between the loss of these two secretory lineages and to avoid the nonspecific effects associated with pharmacological injury models, we generated *Atp4b*-DTR and *Gif*-DTR transgenic mouse models (Figure 1D). These models allow for the selective ablation of parietal and chief cells, respectively, via diphtheria toxin (DT) administration. Efficient and specific cell ablation was confirmed by histology and immunostaining (Figure 1D-E). Interestingly, H/K-ATPase+ parietal cells were significantly reduced even in chief cell-ablated *Gif*-DTR mice. In contrast, parietal cell ablation did not affect the number of GIF+ chief cells, although the proportion of GIF+GSII+ intermediate cells was markedly increased within the GIF+ population. In both *Gif*-DTR and *Gif*-DTR; *Atp4b*-DTR mice, the number of GSII+ neck cells was dramatically elevated. Ki67 staining demonstrated that proliferative activity was predominantly localized to the isthmus and mid-glandular regions, with a significant increase across all DTR mouse models. Moreover, AQP5+ and TFF2+ metaplastic cells emerged following the ablation of either parietal or chief cells, with the highest number observed in mice with combined ablation of both lineages. These results suggest that the loss of a single secretory lineage is sufficient to induce acute SPEM, and that chief cell ablation leads to a reduction in parietal cell numbers.

We generated LSL-*Cdx2* mice, in which the intestine-specific transcription factor Cdx2 can be ectopically expressed in a Cre-dependent manner, in order to investigate the effects by parietal and chief cell loss on intestinal metaplasia (IM) progression. When crossed with *Tff1*-Cre mice, which broadly target the gastric epithelium, Cdx2 was expressed in the upper part of corpus glands without significant histological changes indicative of IM (Figure 1F). When parietal and chief cell ablation was induced using DTR mouse models, parietal cell ablation alone did not alter Cdx2+ cell numbers. However, chief cell ablation, either with or without simultaneous parietal cell ablation, resulted in a dramatic expansion of Cdx2+ cells, accompanied by parietal cell loss (Figure 1F and S1G). In all DTR models, CD44+ metaplastic cells emerged at the base of the glands, with an expansion of the proliferative zone observed above it. The increase in cellular proliferation was most pronounced when both parietal and chief cells were ablated simultaneously. In chief cell-ablated mice, MUC2+Alcian blue+ intestinal goblet-like cells appeared and expanded. Together, these findings demonstrate that chief cell loss not only precedes but also directly contributes to parietal cell loss, driving the progression of gastric atrophy and metaplasia.

### Isthmal neck progenitors are activated and expand in response to acute injury

To investigate alterations in cellular dynamics during mucosal injury and repair, we performed single-cell RNA sequencing (scRNA-seq) analysis on wild-type (WT) mice, WT mice treated with a high dose of tamoxifen as a model of pharmacologically induced acute SPEM^10^ (hereafter referred to as TAM mice), *Atp4b*-DTR (ATP) mice, *Gif*-DTR (GIF) mice, and *Gif*-DTR; *Atp4b*-DTR (GIFATP) mice (Figure 2A). A total of 26,578 cells from the gastric corpus of these mice were classified into 14 clusters based on differentially expressed genes (DEGs) specific to each cluster, including chief cells, neck cells, myeloid cells, myocytes, stem cells, pit cells-1 and -2, parietal cells, fibroblasts-1 and -2, endothelial cells, neuroendocrine cells, lymphoid cells, and nerve cells (Figure 2A and S2A-B).

**Figure 2.**
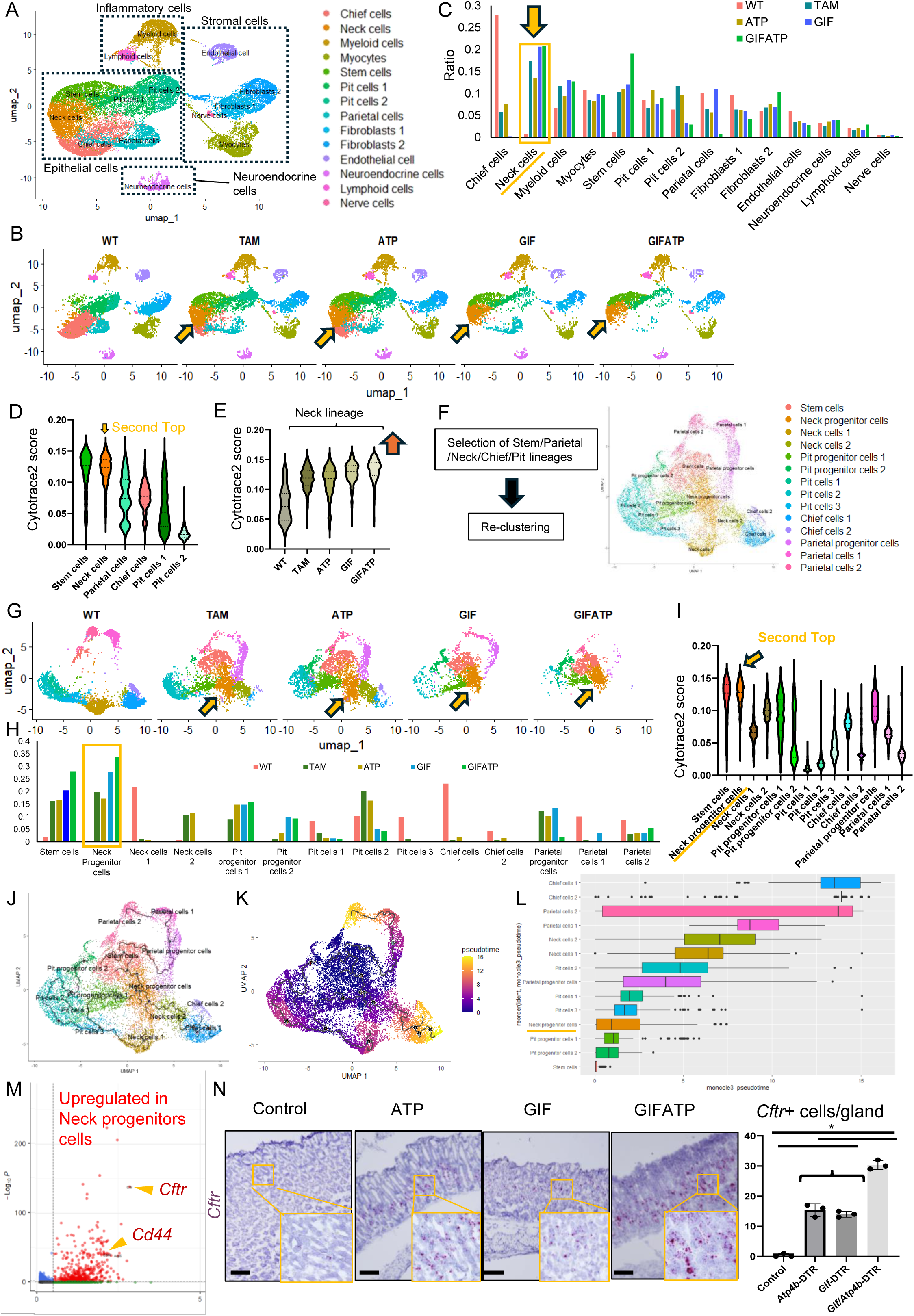
Isthmal neck progenitors are activated and expand in response to acute injury. (A) UMAP plot displaying 26,578 cells from the gastric corpus across 14 clusters. (B) UMAP plot from WT, TAM, ATP, GIF, and GIFATP mice separately visualized. (C) Distribution of cells across the 14 clusters in WT, TAM, ATP, GIF, and GIFATP mice. (D) Violin plots showing Cytotrace2 scores across epithelial cell clusters. (E) Violin plots showing Cytotrace2 scores of the Neck cell cluster in WT, TAM, ATP, GIF, and GIFATP mice. (F) UMAP plot of the gastric epithelial cell clusters after re-clustering. (G) UMAP plot from WT, TAM, ATP, GIF, and GIFATP mice separately visualized. (H) Distribution of cells across the 14 clusters in WT, TAM, ATP, GIF, and GIFATP mice. (I) Violin plots showing Cytotrace2 scores across epithelial cell clusters. (J) Trajectory analysis as visualized in the UMAP plot. (K) Pseudotime calculation visualized in the UMAP plot. (L) Ordering of cell clusters based on Monocle3 pseudotime analysis. (M) Volcano plot displaying the DEGs in the neck progenitor cells. (N) RNA-ISH with *Cftr* probe in WT, ATP, GIF, and GIFATP mice (n=3 mice/group). Scale bars; 100 μm. Mean ± S.E.M. *P < .05.

Notably, the neck cell lineage was significantly expanded in TAM, ATP, GIF, and GIFATP mice compared to WT mice, consistent with histological analyses (Figure 2B-C). Among pit, chief, parietal, neck, and stem cells, stem cells exhibited the highest CytoTRACE2 score^26^, followed by neck cells (Figure 2D). Within the neck cell clusters, samples from TAM, ATP, GIF, and GIFATP mice exhibited higher CytoTRACE2 scores than those from WT mice (Figure 2E). These findings suggest that acute mucosal injury enhances stemness activity within the neck cell lineage.

Next, we conducted an in-depth analysis of epithelial cells—including pit, chief, parietal, neck, and stem cells—followed by re-clustering (Figure 2F). A total of 14,901 cells were newly classified into 14 distinct clusters based on their respective cell markers: stem cells, neck progenitor cells, neck cells-1 and -2, pit progenitor cells-1 and -2, pit cells-1 to -3, chief cells-1 and -2, parietal progenitor cells, and parietal cells-1 and -2 (Figure S2C-D). In TAM, ATP, GIF, and GIFATP mice, we observed an increase in neck progenitor cells and pit/parietal progenitor cells, accompanied by a decrease in neck cells-1, pit cells, chief cells, and parietal cells (Figure 2G). According to the CytoTRACE2 score, neck progenitor cells exhibited the second highest score after stem cells (Figure 2H). Moreover, CytoTRACE2 scores were elevated in stem cells, neck progenitor cells, pit progenitor cells, and parietal progenitor cells following injury, whereas no significant changes were observed in chief cell and other mature cell clusters (Figure S2E). These findings suggest that chief cell de-differentiation does not contribute to mucosal regeneration in this context. Monocle3 pseudotime analysis further delineated the differentiation trajectories within these epithelial cell populations, indicating that neck progenitor cells are directly derived from stem cells and serve as multipotent progenitors, giving rise to pit, neck, and chief cell lineages (Figure 2I).

To identify specific markers for neck progenitor cells, we analyzed upregulated DEGs in neck progenitor cells compared to other clusters (Figure 2J). Among several DEGs, *Cd44* and *Cftr*, both of which have been reported as SPEM markers²³, were significantly upregulated, suggesting that neck progenitor cells contribute to the formation of metaplastic cells. Feature plots confirmed the elevated expression of *Cd44* and *Cftr* in neck progenitor cells following injury (Figure S2F). We conducted spatial transcriptomic analysis to determine the specific anatomical locations of gene expression alterations. In the selected regions corresponding to the glands, the expression of parietal cell markers (*Atp4b, Atp4a*) was decreased in all DTR models, while the expression of proliferation markers (*Mki67*) and metaplasia markers (*Cd44, Aqp5*) were elevated, consistent with findings from scRNAseq (Figure S2G). In addition, *Cftr* expression was greatly increased in the upper or mid-glandular regions of DTR mice (Figure S2H). RNA *in situ* hybridization (ISH) confirmed that *Cftr* expression is nearly absent in the normal stomach but emerges in the upper or mid-glandular regions following chief and parietal cell ablation (Figure 2K). Thus, the loss of chief and parietal cells leads to the expansion of neck progenitor cells, which begin expressing metaplastic markers and likely contribute to mucosal repair (Figures S2I-J).

### Activated Muc6-expressing progenitors are multipotent and serve as the origin of metaplasia

MUC6 is a well-validated marker of gastric neck cells. Our scRNAseq results from WT mouse corpus demonstrated that *Muc6* was most highly expressed in neck cell clusters, and that low levels of *Muc6* expression were also detected in stem cells and other lineages (Figure S3A). We investigated *Muc6*-dsRED-FlpER mice, which specifically express dsRED reporter and tamoxifen-inducible flippase under the control of the *Muc6* promoter (Figure S3B). Consistent with scRNAseq data, a subset of dsRED+ *Muc6*-expressing cells reside above mature neck cell region visualized by GSII staining, co-expressing with stem cell markers Stmn1 and Ki67 (Figure 3A). This suggests that *Muc6* is expressed in not only mature neck cells but also proliferating stem/progenitors in the isthmus. By combining these *Muc6* reporter mice with DTR models, we found that *Muc6*+dsRED+ cells became more proliferative and metaplastic with modest downward migration towards gland base following parietal and/or chief cell ablation (Figure 3B).

**Figure 3.**
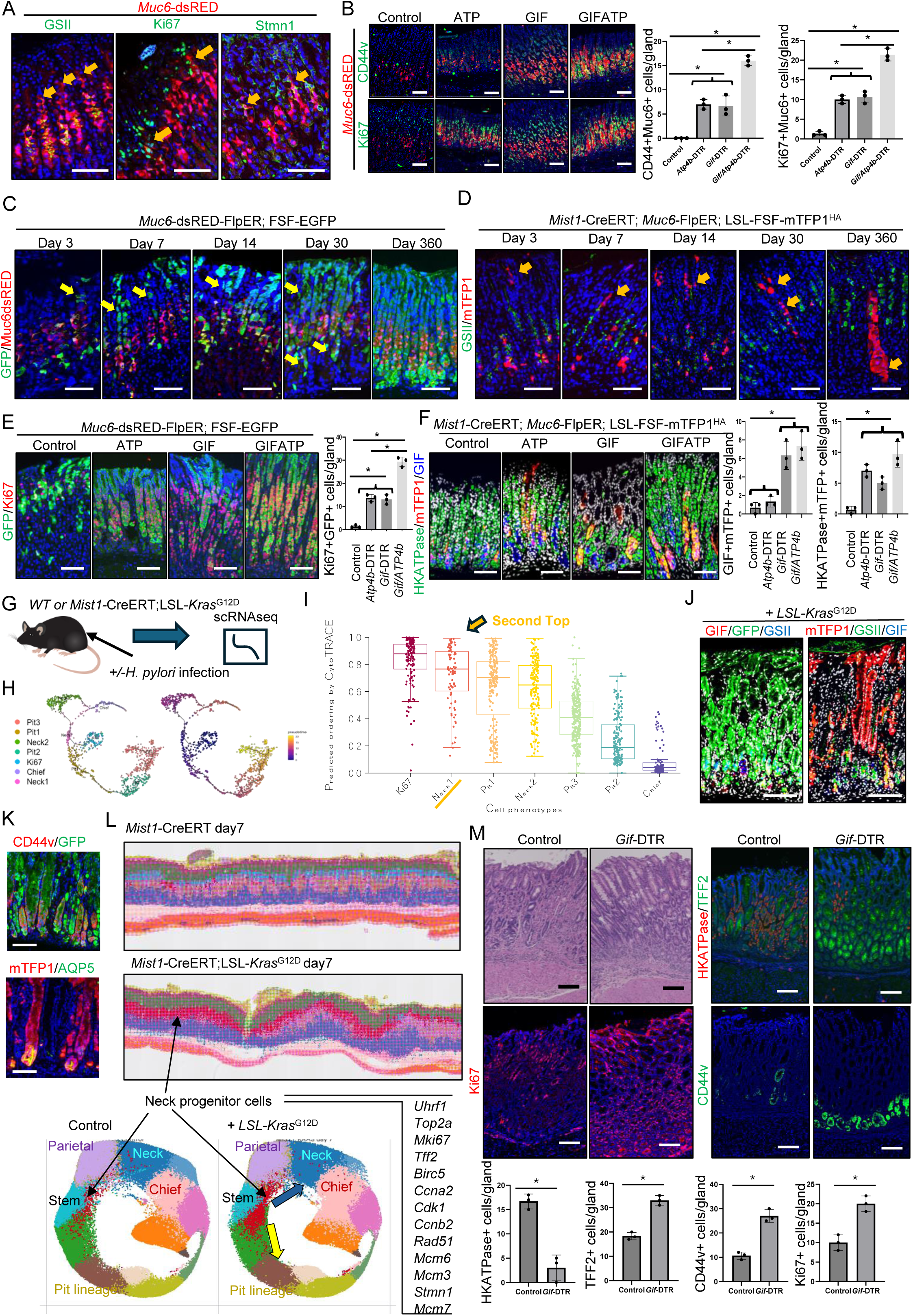
Activated *Muc6*-expressing progenitors are multipotent and serve as the origin of metaplasia. (A) GSII (green)/*Muc6*-dsRED (red), Ki67 (green)/*Muc6*-dsRED (red), and Stmn1 (green)/*Muc6*-dsRED (red) staining in *Muc6*-dsRED-FlpER mice. Arrows indicate GSII-negative, Ki67-positive, or Stmn1-positive *Muc6*+ progenitor cells. (B) Ki67 (green)/*Muc6*-dsRED (red) and CD44v (green)/*Muc6*-dsRED (red) staining in WT, ATP, GIF, and GIFATP mice mated with *Muc6*-dsRED-FlpER mice three days after DT treatment. (C) Lineage tracing images of *Muc6*-dsRED-FlpER; FSF-EGFP mice treated with 300 mg/kg tamoxifen along the time course. (D) Lineage tracing images of *Mist1*-CreERT; *Muc6*-dsRED-FlpER; FSF-LSL-mTFP1^HA^ mice treated with 150 mg/kg tamoxifen along the time course. (E) *Muc6*-dsRED-FlpER; FSF-EGFP mice mated with WT, ATP, GIF, GIFATP mice were treated with 50 mg/kg tamoxifen (day 0). DT was administered at day 3, and GFP(green)/Ki67 (red) expression was stained and quantified at day 10 (n=3 mice/group). (F) *Mist1*-CreERT; *Muc6*-dsRED-FlpER; FSF-LSL-mTFP1^HA^ mice mated with WT, ATP, GIF, GIFATP mice were treated with 150 mg/kg tamoxifen (day 0). DT was administered at day 3, and HKATPase (green)/GIF (blue)/mTFP1 (red) expression was stained and quantified at day 18 (n=3 mice/group). (G-I) Single cell RNA-seq with *Mist1*-CreERT; LSL-*Kras*^G12D^ mice treated with or without *H. pylori.* (G) Schema of the experiment. (H) Trajectory analysis and pseudotime calculation visualized in the UMAP plot. (I) Box plots showing Cytotrace2 scores across epithelial cell clusters. (J-K) *Mist1*-CreERT; LSL-*Kras*^G12D^; *Muc6*-dsRED-FlpER; FSF-EGFP and *Mist1*-CreERT; LSL-*Kras*^G12D^ ; *Muc6*-FlpER; LSL-FSF-mTFP1^HA^ were treated with 150 mg/kg tamoxifen and analyzed at day 30. (J) GIF(red)/GFP(green)/GIF(blue) and mTFP1(red)/GSII(green)/GIF(blue) staining are shown respectively. (K) CD44v(red)/GFP(green) and HA(red)/AQP5(green) staining are shown respectively. (L) Spatial transcriptomic Visium HD analysis of *Mist1*-CreERT; LSL-*Kras*^G12D^ mice 7 days after 150 mg/kg tamoxifen treatment. Spatial gene distribution and UMAP plot split by *Mist1*-CreERT and *Mist1*-CreERT; LSL-*Kras*^G12D^ mice, and DEGs in the neck progenitors are shown. (M) HE, H/K-ATPase (red)/TFF2 (green), RFP (red)/CD44v (green), and Ki67 staining in *Mist1*-CreERT; LSL-*Kras*^G12D^ ; Gif-DTR mice treated with or without DT three times per week for two weeks (n=3 mice/group).

To track the fate of *Muc6*+ lineage cells, we performed lineage tracing experiments using *Muc6*-dsRED-FlpER; FSF-EGFP mice. Shortly after tamoxifen administration, EGFP expression was predominantly induced in the dsRED-high mid-glandular neck cells, whereas occasional EGFP+ cells were observed near the isthmus region, just above dsRED-high cells (Figure 3C). These isthmal EGFP+ clones initially expanded toward the pit cell region near the mucosal surface, followed by downward expansion below the isthmus and dsRED-high regions. A similar bidirectional expansion of EGFP+ cells was observed even after low-dose tamoxifen administration (Figure S3C). Over time, EGFP+ clones progressively occupied entire corpus glands and persisted for over one year, indicating that the *Muc6*+ lineage harbors long-lived multipotent stem or progenitor cells in the corpus.

To more specifically target neck progenitors, we generated *Mist1*-CreERT; *Muc6*-dsRED-FlpER; LSL-FSF-mTfp1^HA^ mice, in which mTFP1 expression is induced in cells co-expressing *Mist1* and *Muc6* (Figure S3D). As previously reported^4^, *Mist1* is expressed in both isthmus stem cells and basal chief cells, and we confirmed co-expression of these genes in a subset of progenitor and chief cells (Figure S3A). Consistently, shortly after tamoxifen induction, mTFP1 expression was detected near the isthmus (corresponding to progenitors) and just above the basal chief cells (corresponding to intermediate cells between GSII+ neck cells and GIF+ chief cells) (Figure 3D). Some mTFP1+ isthmal clones rapidly expanded toward the surface, while others gradually expanded toward the gland base, predominantly tracing neck lineage cells. The mTFP1+ clones migrating downward eventually gave rise to basal chief cells, and a few clones persisted for over one year. Thus, this mTFP1-based labeling system enables targeting of isthmal neck progenitors that give rise to mature neck cells and subsequently to basal chief cells.

To assess the response of these progenitor populations to cellular injury, we ablated parietal and chief cells by combining *Atp4b*-DTR and *Gif*-DTR mice with lineage tracing models. In *Muc6*-dsRED-FlpER; FSF-EGFP mice, we observed a significant increase in Ki67+ proliferating cells within EGFP+ *Muc6*-derived clones following cell ablation (Figure 3E). Similarly, in the mTFP1-based lineage tracing model, ablation of parietal and chief cells accelerated the expansion of neck progenitor-derived lineages with mTFP1+ labeling of newly regenerated parietal and chief cells (Figure 3F). These findings suggest that *Mist1*+*Muc6*+ isthmal neck progenitors respond to the loss of parietal and chief cells and contribute to mucosal regeneration.

Previous studies have demonstrated that oncogenic Kras activation leads to the development of metaplasia in the stomach^4^. To further elucidate the cellular dynamics underlying Kras-dependent metaplasia development, we analyzed scRNA-seq data from gastric epithelial cells of *Mist1*-CreERT; LSL-*Kras*^G12D^ mice^27^ (Figure 3G). UMAP clustering identified major epithelial cell populations, with key markers confirming cluster identities (Figure S3E). Comparative analysis between groups revealed notable alterations in epithelial cell distribution, indicating that the neck-2 cluster contained Kras-induced metaplastic cells (Figure S3F). CytoTRACE2 analysis demonstrated significant variations across epithelial cell clusters, with neck-1 cells exhibiting the second highest score after *Stmn1*+*Mki67*+ stem cells, consistent with our DTR-mice data (Figure 3H-I). Trajectory inference and pseudotime analyses indicated a dynamic transition from the neck-1 cluster to the neck-2 metaplastic cluster, suggesting that the neck-1 cluster likely represents neck progenitors that give rise to metaplasia. Indeed, we confirmed in lineage tracing experiments that *Muc6*-derived EGFP+ and mTFP1+ clones expanded more rapidly following mutant Kras induction in *Mist1*+ cells, eventually dominating entire corpus glands with robust expression of metaplasia markers CD44 and AQP5 (Figure 3J-K).

To determine where Kras-driven metaplasia arises from, we performed spatial transcriptomic analysis in *Mist1*-CreERT; LSL-*Kras*^G12D^ mice shortly after tamoxifen induction to identify the spatial organization. Compared to *Kras*-intact controls, mutant *Kras* induction in the *Mist1*+ lineage led to bidirectional expansion of isthmal progenitor cells expressing stem cell and neck lineage markers (Figure 3L and S3G-H). These transcriptomic analyses strongly indicate that *Kras*-induced metaplasia in *Mist1*-CreERT mice originates from isthmal neck progenitors. Similar to findings in the Cdx2-dependent IM model, additional chief cell ablation using *Gif*-DTR mice further accelerated metaplasia progression in *Mist1*-CreERT; LSL- *Kras*^G12D^ mice (Figure 3M), reinforcing the idea that Kras-induced metaplasia does not originate from GIF+ chief cells but rather from isthmal progenitors. Additionally, we confirmed that metaplastic glands in *Helicobacter pylori*-infected mice were efficiently traced by the *Muc6*+ lineage. Collectively, these findings indicate that *Muc6*+ isthmal progenitors serve as the primary origin of various types of metaplasia in the stomach, under both injury-induced and pro-oncogenic conditions.

### NRG1-producing fibroblasts activate Myc Signaling in isthmus progenitors and serve as a metaplastic niche following injury

To explore global gene expression changes that occurred after loss of parietal and/or chief cells, we performed bulk RNA-seq analysis of the gastric corpus from WT, ATP, GIF, and GIFATP mice. Principal component analysis (PCA) revealed a clear separation among the four groups (Figure S4A). A volcano plot, Venn diagram, and heatmap identified significant upregulation of stem cell markers (*Mki67*, *Stmn1*) and metaplasia markers (*Aqp5*, *Tff2*, *Cd44*, etc) in DTR mice compared to control mice (Figure S4B-D and Table S1-3). Among the 1,260 upregulated differentially expressed genes (DEGs) and 354 downregulated DEGs, genes typically expressed in normal gastric cell lineages—including *Atp4a*, *Atp4b*, and *Gif*—were significantly downregulated. Gene set enrichment analysis (GSEA) using hallmark gene sets revealed that DTR-treated samples were particularly enriched for genes related to mitosis and cell cycle regulation, including Myc, E2F, and G2M checkpoint signaling (Figure 4A and Table S4).

**Figure 4.**
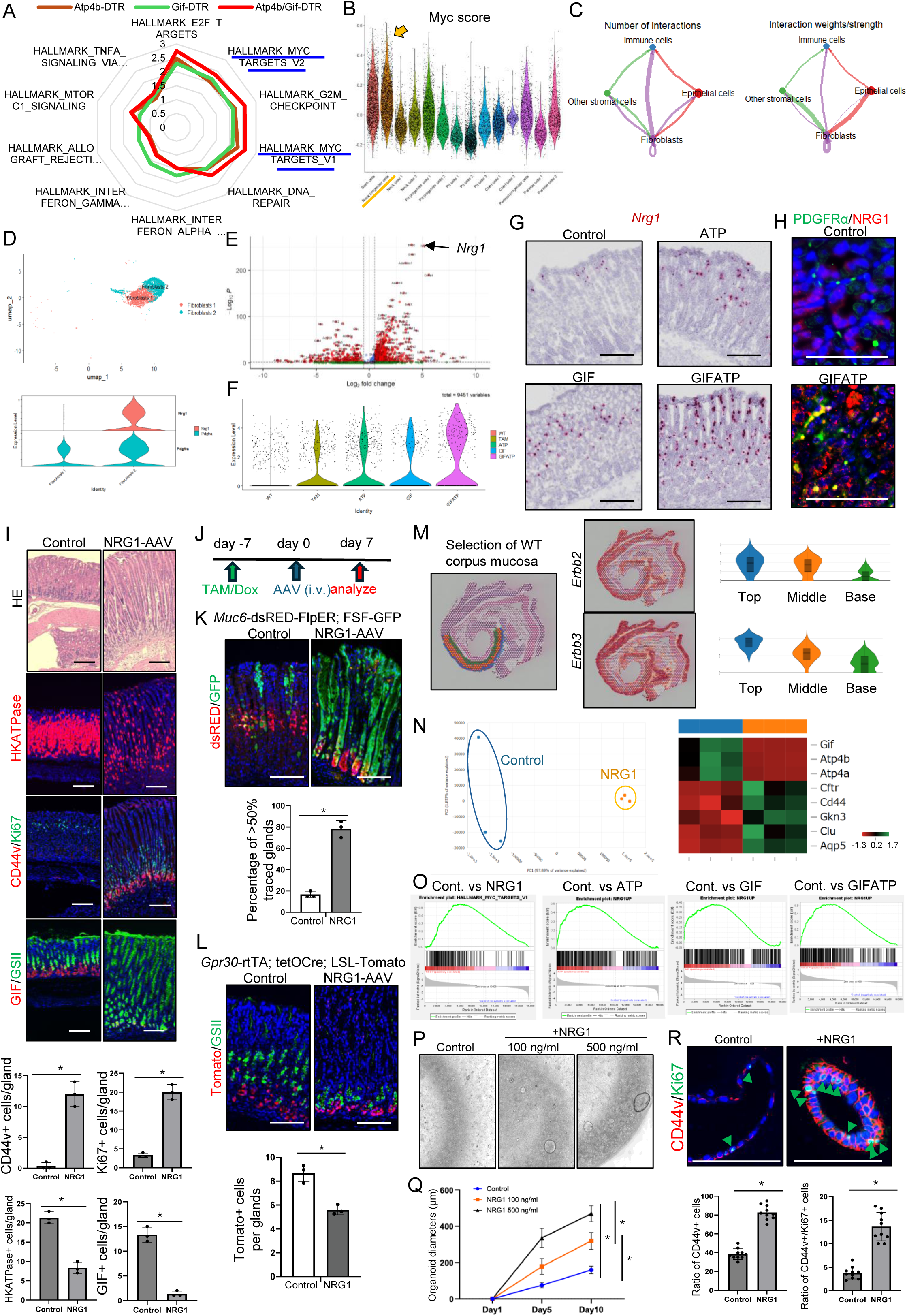
NRG1-producing fibroblasts activate Myc Signaling in isthmus progenitors and serve as a metaplastic niche following injury. (A) Radar chart showing the GSEA analysis of bulk RNA seq with Hallmark gene sets. (B) Myc score across gastric epithelial cell clusters of scRNA seq is shown in a violin plot. (C) Circle plot showing the cell-to-cell interaction reflecting the number of interactions and interaction wright/strength. (D) UMAP plot and violin plot of *Nrg1 and Pdgfra* expression in the fibroblast clusters. (E) Volcano plot displaying the DEGs of the fibroblasts 2 cluster compared with the fibroblast 1 cluster. (F) Violin plot of *Nrg1* expression in WT, TAM, ATP, GIF, and GIFATP mice. (G) RNA-ISH with *Nrg1* probe in WT, ATP, GIF, and GIFATP mice. (H) PDGFRa (green)/NRG1 (red) staining in WT and GIFATP mice. (I) HE, H/K-ATPase, CD44v (green)/Ki67 (red), and GSII (green)/GIF (red) staining and quantification of mice treated with control and NRG1-AAV seven days after injection (n=3 mice/group). (J-L) Lineage tracing experiments with NRG1-AAV treatment. (J) Schema of the experiments. (K) *Muc6*-dsRED-FlpER; FSF-EGFP mice were treated with 150 mg/kg tamoxifen 7 days prior to AAV administration. GFP (green)/*Muc6*-dsRED (red) staining and percentages of >50% tracing events are shown. (L) *Gpr30*-rtTA; tetOCre; LSL-Tomato mice were treated with 0.2% doxycycline for 3 days. Tomato (red)/GSII (green) staining and numbers of Tomato+ cells are shown. (M) Spatial gene distribution of *Erbb2* and *Erbb3* and violin plot across top/middle/base corpus glands of WT mice. (N) PCA plot and heatmap showing *Gif, Atp4b, Atp4a, Cftr, Cd44, Gkn3, Clu, Aqp5* expression comparing the bulk RNA seq between control and NRG1-AAV treated mice. (O) The result of GSEA analysis. Enrichment of Myc target genes in NRG1-AAV treated mice compared to control mice, and enrichment of upregulated DEGs after NRG1-AAV treatment in ATP, GIF, and GIFATP mice compared to WT mice are shown as sigmoid curves. (P-Q) Proliferation assay with mice gastric organoids treated with or without recombinant NRG1. (R) CD44v (green)/Ki67 (red) staining in control and NRG1-treated mice organoids.

Based on our scRNA-seq data, Myc signaling was most highly activated in the stem and neck progenitor clusters (Figure 4B and S4E-F). Indeed, c-Myc phosphorylation was significantly increased in the mid-glandular region of injured corpus glands (Figure S4G). Given that stem and progenitor activation is regulated by the surrounding stromal niche, we performed cell-cell interaction analysis, which identified fibroblasts as a central hub of intercellular communication, with strong outgoing signaling toward the stem and neck progenitor clusters (Figure 4C and S4H). Our scRNA-seq analysis of acute injury mouse models identified two distinct fibroblast subpopulations (Figure 4D). Differential gene expression (DEG) analysis identified key fibroblast markers within each population and demonstrated that *Nrg1* was the most prominently upregulated gene in the fibroblast-2 cluster (Figure S2B). *Nrg1* expression was significantly higher in the fibroblast-2 cluster compared to the fibroblast-1 cluster and was further increased in TAM, ATP, GIF, and GIFATP mice compared to WT controls (Figure 4D-F and S4I-J). ISH demonstrated that *Nrg1* expression is localized to stromal cells near the isthmus in normal stomach tissue and is upregulated in injury models (Figure 4G). Co-expression analysis of *Nrg1* and *Pdgfra* further confirmed that fibroblasts are the primary source of NRG1 production, and this was validated by IHC (Figure 4D and 4H).

Previous studies suggest that NRG1 plays a crucial role in intestinal mucosal healing^28^ and that NRG1 signaling is upregulated in human gastric metaplasia^29^. To assess the functional role of NRG1 in the stomach, we administered an NRG1-expressing AAV vector to mice (Figure 4I). NRG1 overexpression induced marked mucosal hypertrophy, with an increase in CD44v+ metaplastic cells and Ki67+ proliferative cells, along with a decrease in GIF+ chief cells and H/K-ATPase+ parietal cells. These histological and molecular changes closely resembled those observed during mucosal regeneration and metaplasia progression. Among the EGF family members, we realized that *Areg* was also upregulated in injury models besides *Nrg1*, particularly within the pit cell clusters (Figure S4K). Nevertheless, when we administered an AREG-expressing AAV vector to mice, no major histological changes were observed (Figure S4L), suggesting that NRG1, rather than AREG, likely acts as a primary metaplastic niche factor near the isthmus stem/progenitor cells.

During NRG1 overexpression, *Muc6*-derived lineage tracing demonstrated a marked expansion of isthmus-derived EGFP+ clones, with a dramatic downward shift of *Muc6*+dsRED+ progenitor cells (Figure 4J-K). In contrast, *Gpr30*-expressing chief cells did not undergo upward expansion and instead showed a significant reduction (Figure 4L), validating that NRG1-driven hyperplastic changes originate from isthmus progenitors, not from basal chief cells. Indeed, NRG1 receptors, ERBB2 and ERBB3, were expressed in gastric epithelial cells, particularly within the pit, stem, and neck clusters, as shown by single-cell RNA-seq and spatial transcriptomic analysis (Figure 4M and S4M).

Bulk RNA-seq analysis of NRG1-AAV-treated mice revealed upregulation of metaplastic markers and activation of Myc signaling, with a significant increase in phospho-c-Myc+ cells in NRG1-overexpressing mice (Figure 4N-O and S4N). Notably, the gene expression signatures in DTR-treated mice showed a strong correlation with NRG1-mediated gene alterations, as demonstrated by GSEA (Figure 4O). To further validate NRG1’s role in epithelial remodeling, we treated gastric organoids with recombinant NRG1, which resulted in increased proliferation and upregulation of metaplasia markers such as CD44v (Figure 4P-R). Taken together, these findings indicate that NRG1 secreted by stromal cells plays a critical role in mucosal regeneration and metaplasia development following injury, primarily through the activation of Myc signaling.

### Monocyte-derived IL1β controls mucosal regeneration and metaplasia development by promoting NRG1 secretion from fibroblasts

To investigate the mechanism underlying NRG1 upregulation in fibroblasts during gastric injury, we hypothesized that inflammatory signaling plays a key role, given the importance of immune-stromal crosstalk in various biological responses. We re-clustered immune cell populations from the acute injury scRNA-seq dataset and identified ten distinct immune subclusters (Figures 5A and S5A-B). Among them, the inflammatory monocyte cluster was significantly increased following tissue injury based on CIBERSORTx analysis^30^ of bulk RNA-seq data (Figure S5C). The inflammatory monocyte cluster was enriched for proinflammatory genes, including *Il1a* and *Il1b*, whose expression was further elevated in TAM, ATP, GIF, and GIFATP mice (Figure 5B). DEGs in inflammatory monocytes also included *Msr1*, a macrophage/monocyte marker (Figures 5C-D). IHC analysis revealed increased infiltration of MSR1+ cells in ATP, GIF, and GIFATP mice (Figure 5E). To assess the role of MSR1+ inflammatory monocytes, we subjected *Msr1*-DTR mice to an HDT-induced injury model (Figures 5F). DT treatment in *Msr1*-DTR mice successfully depleted MSR1+ monocytes, leading to a significant reduction in H/K-ATPase+ parietal cells and GIF+ chief cells, suggesting that MSR1+ monocytes are essential for the regeneration of these lineages following acute injury (Figure 5G-H).

**Figure 5.**
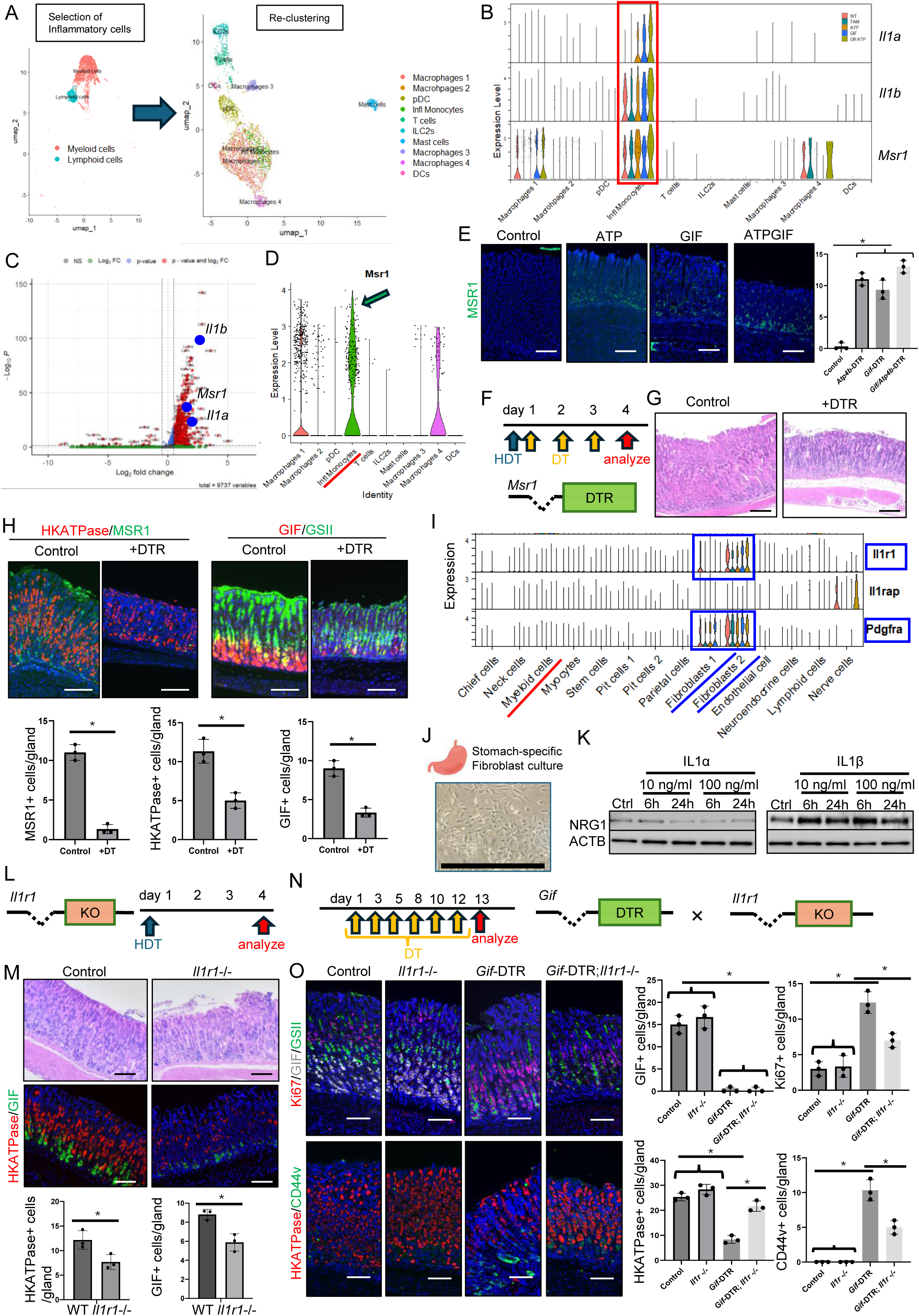
Monocyte-derived IL1β controls mucosal regeneration and metaplasia development by promoting NRG1 secretion from fibroblasts. (A) Re-clustering of the gastric inflammatory cell clusters. (B) Violin plot of *Il1a, Il1b*, and *Msr1* expression across inflammatory cell clusters. (C) Volcano plot displaying the DEGs in the inflammatory monocytes cluster. (D) Violin plot of *Msr1* expression across 10 inflammatory cell clusters. (E) MSR1 staning in WT, ATP, GIF, and GIFATP mice. (F-H) *Msr1*-DTR mice experiments with high-dose (300 mg/kg) tamoxifen treatment. (F) Protocol and gene constructs. (G) HE and (H) MSR1 (green)/H/K-ATPase (red), and GSII (green)/GIF (red) staining in control and DT-treated mice (n=3 mice/group). (I) Violin plot of *Il1r1, Il1rap, and Pdgfra* expression across all clusters. (J) Microscopic image of the cultured fibroblasts isolated from mouse corpus. (K) IL1α and IL1β administration in the gastric fibroblast culture. Western blots for NRG1 and β-actin are shown. (L-M) *Il1r1-/-* mice were treated with high-dose (300 mg/kg) tamoxifen. (L) Protocol and gene constructs. (M) HE and GIF (green)/H/K-ATPase (red) staining in control and *Il1r1-/-* mice (n=3 mice/group). (N-O) *Il1r1-/-; Gif*-DTR mice experiments. (N) Protocol and gene constructs. (O) GSII (green)/Ki67 (red)/GIF (gray) and H/K-ATPase (red)/CD44v (green) staining in control, *Il1r1*-/-, *Gif*-DTR, and *Gif*-DTR; *Il1r1*-/- mice.

We next examined the expression of *Il1a, Il1b*, and their main receptor *Il1r1* in the total acute injury scRNA-seq dataset. While *Il1a* and *Il1b* were predominantly expressed in *Ptprc+* immune cells, *Il1r1* was preferentially expressed in *Pdgfra*+ fibroblasts, suggesting the presence of an IL1-mediated immune-stromal signaling network (Figure 5I and S5D). To confirm this, we isolated gastric fibroblasts from mouse stomachs and treated them *ex vivo* with IL1α or IL1β (Figure 5J). Following treatment, IL1β, but not IL1α, significantly increased NRG1 expression in fibroblasts, indicating that IL1β serves as the primary mediator of this immune-stromal interaction (Figure 5K).

To determine whether IL1 signaling is required for acute regeneration, we treated *Il1r1*⁻/⁻ mice with HDT (Figure 5L). Similar to *Msr1*-DTR mice, *Il1r1*⁻/⁻ mice exhibited a significant reduction in parietal and chief cell numbers compared to WT controls shortly after HDT treatment (Figure 5M), suggesting that IL1R1+ fibroblasts, activated by monocyte-derived IL1β, promote the regeneration of these two lineages following acute injury. Next, we investigated the role of IL1 signaling in chronic gastric injury by administering continuous DT treatment to *Gif-*DTR mice, a model that induces sustained chief cell loss followed by progressive parietal cell loss (Figures 5N). After two weeks of DT treatment, *Il1r1* knockout in *Gif*-DTR mice resulted in a reduction in CD44v+ metaplastic cells and Ki67+ proliferating cells while preserving H/K-ATPase+ parietal cells (Figure 5O). These findings suggest that while IL1 signaling and subsequent NRG1 upregulation are required for the acute regeneration of secretory lineages, blocking IL1 signaling in the chronic phase prevents atrophic and metaplastic changes. Together, IL1R1/NRG1 signaling plays a fundamental role in gastric epithelial remodeling via immune-stromal interactions.

### IL1R1/NRG1-dependent interaction is present in the human atrophic and metaplastic stomach

To assess the clinical relevance of our findings in mice, we analyzed human gastric samples from patients with atrophic gastritis. Immunofluorescence staining revealed a progressive increase in NRG1 expression in stromal cells, which correlated with the severity of gastric atrophy (Figure 6A). Next we examined the relationship between mononuclear cell infiltration and atrophy grade. A positive correlation was observed between these parameters, suggesting that mononuclear cell accumulation is associated with gastric epithelial remodeling (Figure 6B).

**Figure 6.**
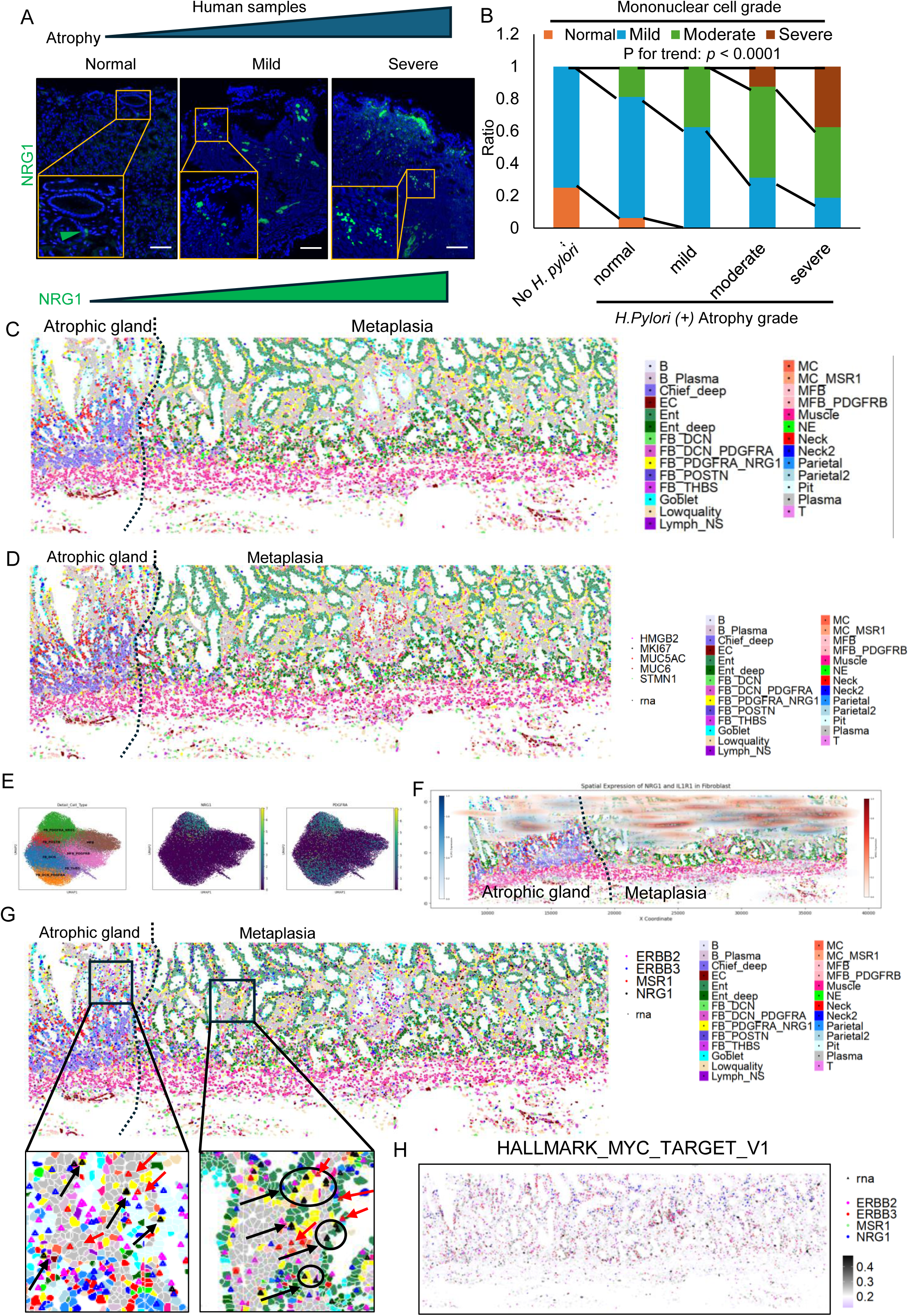
IL1R1/NRG1-dependent interaction is present in human atrophic and metaplastic stomach. (A) NRG1 (red) staining in human samples of atrophic gastritis. (B) The relationship between atrophy grade and mononuclear cell grade. (C-G) Spatial transcriptomic analysis in human gastritis samples. The specimen contains the atrophic region (left) and metaplastic region (right). (C) Cell type annotation in the spatial coordinates. (D) Expression of *HMGB2, MKI67, MUC5AC, MUC6,* and *STMN1* is shown with cell type annotation. (E) UMAP plots of stromal fibroblast compartments. (left) Subcluster annotation. (central) *NRG1* expression. (right) *PDGFRA* expression. (F) *NRG1* (red) and *IL1R1* (blue) kernel density plot showing expression derived exclusively from fibroblasts. (G) Expression of *MSR1, NRG1, ERBB2,* and *ERBB3* is shown with cell type annotation. (H) GSEA scores of Hallmark Myc target V1. In the enlarged views, the color of each arrow corresponds to the respective genes. Abbreviations: EC: Endothelial cell; Ent: Enterocyte; FB: Fibroblast; Lymph_NS: Lymphocyte, non-specific; MC: Myeloid cell; MFB: Myofibroblast; NE: Neuroendocrine cell

To further validate these molecular changes in human samples, we analyzed the spatial transcriptomic data using CosMx 6K panel, adding specific gastric marker genes. A total of 439,454 cells were obtained and subsequently clustered as described in the Methods section. In brief, fibroblast subtypes were annotated based on subtype-specific marker genes, while other cell types were annotated using known lineage markers (Figure 6C). We confirmed the distribution of MUC6+ epithelial cells in the isthmus region together with proliferative marker genes MKI67, HMGB2, and STMN1 (Figure 6D). NRG1 expression in fibroblasts was significantly enriched in PDGFRA+ fibroblasts (Figure 6E), as shown in the UMAP representation. Furthermore, spatial mapping revealed NRG1 expression—visualized selectively in fibroblasts—was more evident in the isthmus zone of both atrophic and metaplastic regions, most of which was co-expressed with IL1R1 (Figure 6F). Scattered accumulation of *MSR1* and *IL1B* expression, shown specifically in myeloid cells, was also confirmed, particularly near the NRG1-expressing fibroblast region (Figure S6A). CD44, displayed only from epithelial cells was detected in the isthmus of atrophic glands, and more robustly expressed in the basal metaplastic region (Figure S6B). There was a moderate proximity between NRG1+ fibroblasts and neck cells in fundic gland regions adjacent to intestinal metaplasia (Figure S6C), coincided with upregulation of ERBB2 and ERBB3 as well as Myc activation in neighborhood epithelial compartments (Figure 6G-H). These findings indicate that IL1R/NRG1-dependent metaplastic niche is present in human tissues and linked to gastric atrophy and metaplasia progression.

## Discussion

Our study provides novel insights into the cellular and molecular mechanisms driving gastric atrophy and metaplasia, demonstrating that chief cell loss precedes and actively contributes to parietal cell depletion in human gastric samples. This finding challenges the conventional paradigm that parietal cell loss is the primary initiating event in gastric mucosal transformation. Instead, our results highlight a previously unrecognized role of chief cells in maintaining gastric epithelial homeostasis, with their loss acting as a critical early driver of mucosal remodeling and metaplastic progression.

A critical finding of our study is that metaplasia originates from neck progenitor cells rather than chief cells. While previous models suggested that acute injury induces chief cell de-differentiation into metaplastic cells, our data indicate that chief cell loss itself drives the expansion of *Mist1*□*Muc6*□ isthmal neck progenitors, which serve as the true source of metaplastic cells, consistent with previous findings^17,18^. This is further supported by single-cell and spatial transcriptomic analysis, which identified an expansion of the neck cell lineage with increased stemness properties and the capacity for metaplastic differentiation particularly after loss of chief cells. Therefore, targeting chief cells for prevention of gastric neoplasia seems to be an erroneous strategy, as it may cause activation of neck lineage that leads to metaplasia progression. Notably, spasmolytic polypeptide (SP/TFF2)—a defining marker of SPEM—is initially expressed in neck cells^3,19^, reinforcing the idea that metaplasia arises from the reprogramming of neck progenitors rather than chief cell de-differentiation. This fundamental shift in our understanding of gastric mucosal remodeling suggests that targeting neck progenitor cells could provide a novel therapeutic strategy to prevent gastric metaplasia and its progression to neoplasia.

Emerging evidence underscores the pivotal role of the c-Myc oncogene in gastric tumorigenesis and metaplastic progression. Myc overexpression and activation have been frequently observed in gastric cancer and intestinal metaplasia, suggesting its role in epithelial reprogramming and tumor initiation^31–34^. Our transcriptomic analysis revealed that Myc signaling is significantly upregulated in metaplastic lesions, primarily mediated by NRG1-dependent pathway. Previous studies demonstrate that transcriptomic and epigenetic analyses in gastric tissues reveal that Myc-driven reprogramming cooperates with inflammatory signaling to disrupt gastric lineage identity, thereby facilitating intestinal differentiation and metaplastic transformation. Given its frequent dysregulation in metaplastic and cancerous gastric tissues, future studies should explore whether targeting Myc-dependent transcriptional programs could serve as a therapeutic strategy to prevent metaplasia progression and gastric tumorigenesis.

Beyond epithelial cell-intrinsic mechanisms, our study underscores the pivotal role of stromal-epithelial interactions in gastric remodeling. Cell-cell interaction analysis identified PDGFRA□ fibroblasts as key mediators of the injury response, with these stromal cells exhibiting upregulated NRG1 expression in an IL1R-dependent manner. Importantly, our findings align with previous research in the intestinal epithelium, where NRG1 has been shown to drive intestinal stem cell proliferation and enhance epithelial regeneration^35,36^. These studies demonstrated that mesenchymal-derived NRG1 promotes epithelial renewal under both homeostatic and injury conditions, a mechanism that appears to be conserved in the gastric mucosa. However, the role of IL1β in regulating NRG1 expression appears to be context-dependent. While our study found that monocyte-derived IL1β stimulates NRG1 secretion from fibroblasts, facilitating epithelial remodeling, a contrasting study in colitis suggests that IL1β suppresses NRG1 expression, delaying epithelial repair^37^. This discrepancy may reflect differences in tissue-specific immune microenvironments or the distinct regulatory networks governing regeneration in the stomach and intestine.

Importantly, our findings indicate that IL1β is not only a key driver of metaplasia but also plays a well-established role in gastric tumorigenesis. Previous studies have demonstrated the potential link between IL1β alterations and gastric inflammation and cancer^38–40^. Notably, in mouse models, IL1β overexpression induces spontaneous gastric inflammation and neoplasia, while deletion of *Il1r1* suppresses *Helicobacter* infection-induced metaplasia development, suggesting that its role in metaplasia may represent an early step in gastric carcinogenesis. Our transcriptomic data in human samples support a pathogenic link between inflammatory cytokine signaling, stromal reprogramming, and gastric epithelial transformation, reinforcing the importance of IL1β/NRG1 signaling in both gastric metaplasia and tumor progression. Future studies should explore whether therapeutic intervention in these pathways can mitigate gastric disease progression and reduce the risk of malignant transformation.

## Author contributions

Junya Arai, MD, PhD (conceptualization: supporting; data curation: Lead; Formal analysis: Lead; Investigation: Lead; Methodology: Lead; Validation: Lead; Visualization: Lead; Writing – original draft: lead.) Yusuke Iwata, MD, (Writing-Review and Editing: Supporting Information). Ayumu Tsubosaka, MD, (Writing-Review and Editing: Supporting Information). Hiroto Kinoshita, MD, PhD (Formal analysis: Supporting; Writing-Review and Editing: Supporting Information). Shintaro Shinohara, MD (Formal analysis: Supporting; Writing-Review and Editing: Supporting Information). Sohei Abe, MD, PhD (Formal analysis: Supporting; Writing, review, and editing: Supporting Information). Toshiro Shiokawa, MD, (Writing-Review and Editing: Supporting Information). Katsuyuki Oura, MD, (Writing-Review and Editing: Supporting Information). Nobumi Suzuki, MD, PhD (Formal analysis: Supporting; Writing-Review and Editing: Supporting Information). Masahiro Hata, MD, PhD (Data curation: Supporting; Resources: Supporting; Writing – review and editing: Supporting). Ken Kurokawa, MD, PhD (Writing-Review and Editing: Supporting Information). Yukiko Oya, MD, PhD (Writing-Review and Editing: Supporting Information). Mayo Tsuboi, MD, PhD (Writing-Review and Editing: Supporting Information). Sozaburo Ihara, MD, PhD (Writing, Review, and Editing: Supporting Information). Keita Murakami, MD (Formal analysis: supporting; Writing-Review and Editing: Supporting Information). Chihiro Shiomi, MD, (Writing-Review and Editing: Supporting Information). Chie Uekura, MD, (Writing-Review and Editing: Supporting Information). Hiroaki Fujiwara, MD, PhD (Writing – Review and Editing: Support). Hiroaki Tateno, Ph.D (Writing – Review and Editing: Support). Seiya Mizuno, PhD (Resources: Supporting; Writing-Review and Editing: Supporting Information). Satoru Takahashi, MD, PhD (Resources: Supporting; Writing-Review and Editing: Supporting Information). Tetsuo Ushiku, MD, PhD (Writing-Review and Editing: Supporting Information). Yoshihiro Hirata, MD, PhD (Writing-Review and Editing: Supporting Information). Hideaki Ijichi, MD, PhD (Writing-Review and Editing: Supporting Information). Masato Kasuga, MD, PhD (Writing-Review and Editing: Supporting Information). Valerie P. O’Brien, PhD (Writing-Review and Editing: Supporting Information), Nina Salama PhD (Writing-Review and Editing: Supporting Information), Miwako Kakiuchi, MD, PhD (Writing-Review and Editing: Supporting Information). Shunpei Ishikawa, MD, PhD (Writing-Review and Editing: Supporting Information). Timothy C. Wang, MD, PhD (conceptualization: support; supervision: support; writing: original draft: support; writing: review and editing: support). Yoku Hayakawa, MD, PhD (Conceptualization: Lead; Data curation: Lead; Formal analysis: Lead; Funding acquisition: Lead; Investigation: Lead; Methodology: Lead; Supervision: Lead; Validation: Supporting; Visualization: Supporting; Writing – original draft: Lead; Writing – review and editing: lead). Mitsuhiro Fujishiro, MD, PhD (project administration: support; supervision: support; writing: review, and editing: support).

## Acknowledgments

We thank the staff members at the laboratory of Gastroenterology, Graduate School of Medicine, University of Tokyo and The Institute of Medical Science, Asahi Life Foundation, especially Sayaka Ito, Hiromi Kato, and Akiko Kikuchi, for their skilled assistance. We also thank Gen Nishitai and Masato Tanaka for providing us the *Msr1*-DTR mice.

This study was supported by Daiwa securities Foundation, Kato Memorial Bioscience Foundation, UBE Foundation, Mochida Memorial Foundation for Medical and Pharmaceutical Research, Takeda Science Foundation, Kobayashi Foundation for Cancer Research, International Medical Research Foundation, Cell Science Research Foundation (to J.A.), Yakult Bio-Science Foundation, Mochida Memorial Foundation for Medical and Pharmaceutical Research, Yasuda Memorial Foundation, Ono Pharmaceutical Foundation for Oncology, Immunology, and Neurology, Eli Lilly Japan KK Innovation Research Grant 2023, Research Grant of the Princess Takamatsu Cancer Research Fund, Suzuken Memorial Foundation, the Mitsubishi Foundation, Kurata Grants from Hitachi Foundation (to Y.H.), JSPS KAKENHI Grants-in-Aid for Scientific Research (no. 24K18531(J.A.), 23H02744 (Y. Hayakawa), JP22H04925 (PAGS), 22H04990 (S. Ishikawa)), P-CREATE/PROMOTE from AMED (Y. Hayakawa), AMED (no. JP23tk0124002 (S. Ishikawa)), the Advanced Research and Development Programs for Medical Innovation (PRIME) (Y. Hayakawa), and NIH/NCI funding including R01DK48077 and R35CA210088 (T.C.W.) and R00CA263036 (V.P.O.).

## Supplementary Figure and table legends

**Figure S1, related to Figure 1.**
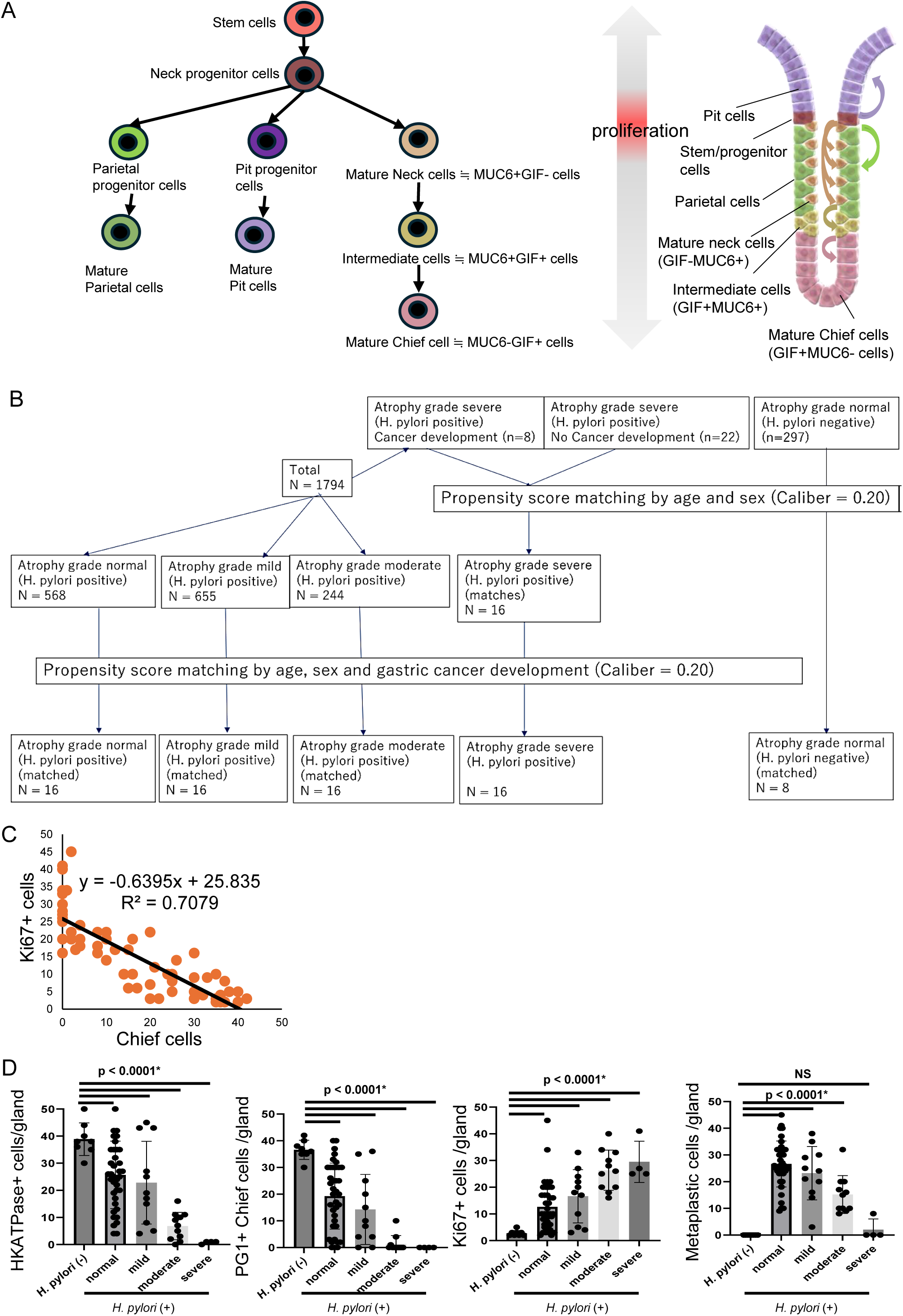
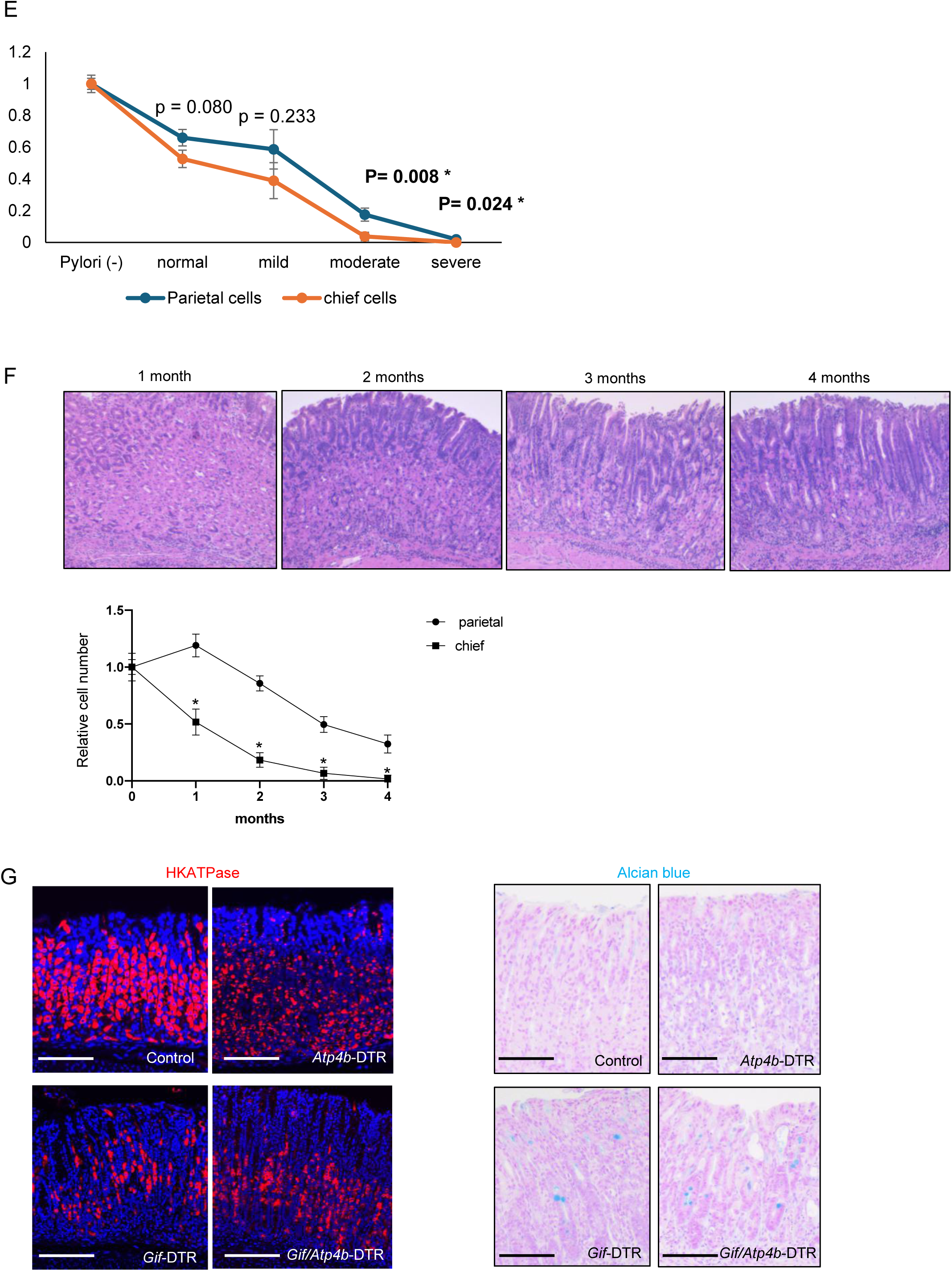
Changes in gastric parietal, chief, and metaplastic lineages during atrophic gastritis. (A) Schema of gastric corpus stem cell hierarchy and gland structure. (B) Patients flow. (C) Comparison of the number of chief cells and Ki67+ cells shown as scatter diagram. (D) H/K-ATPase+ parietal cells, pepsinogen I+ chief cells, Ki67+ proliferative cells, and TFF2+/CD44v+ metaplastic cells per gland according to the intestinal metaplasia grade were quantified. (E) Relative numbers of parietal cells and chief cells according to the severity of Intestinal metaplasia were quantified. Numbers in *H. pylori* (-) group were set as 1.0. (F) HE image of mice infected with *H. pylori* (PMSS1) for 1, 2, 3, and 4 months. Relative numbers of parietal cells and chief cells were compared (n=3 mice/group). Numbers in *H. pylori* (-) group were set as 1.0. (G) HKATPase (red) and Alcian blue staining in *Tff1*-cre; LSL-*Cdx2, Tff1*-cre; LSL-*Cdx2*; *Gif-*DTR, *Tff1*-cre; LSL-*Cdx2*; *Atp4b*-DTR, *Tff1*-cre; LSL-*Cdx2*; *Gif*-DTR; *Atp4b*-DTR mice (n=3 mice/group). Scale bars; 100 μm. Mean ± S.E.M. *P < .05.

**Figure S2, related to Figure 2.**
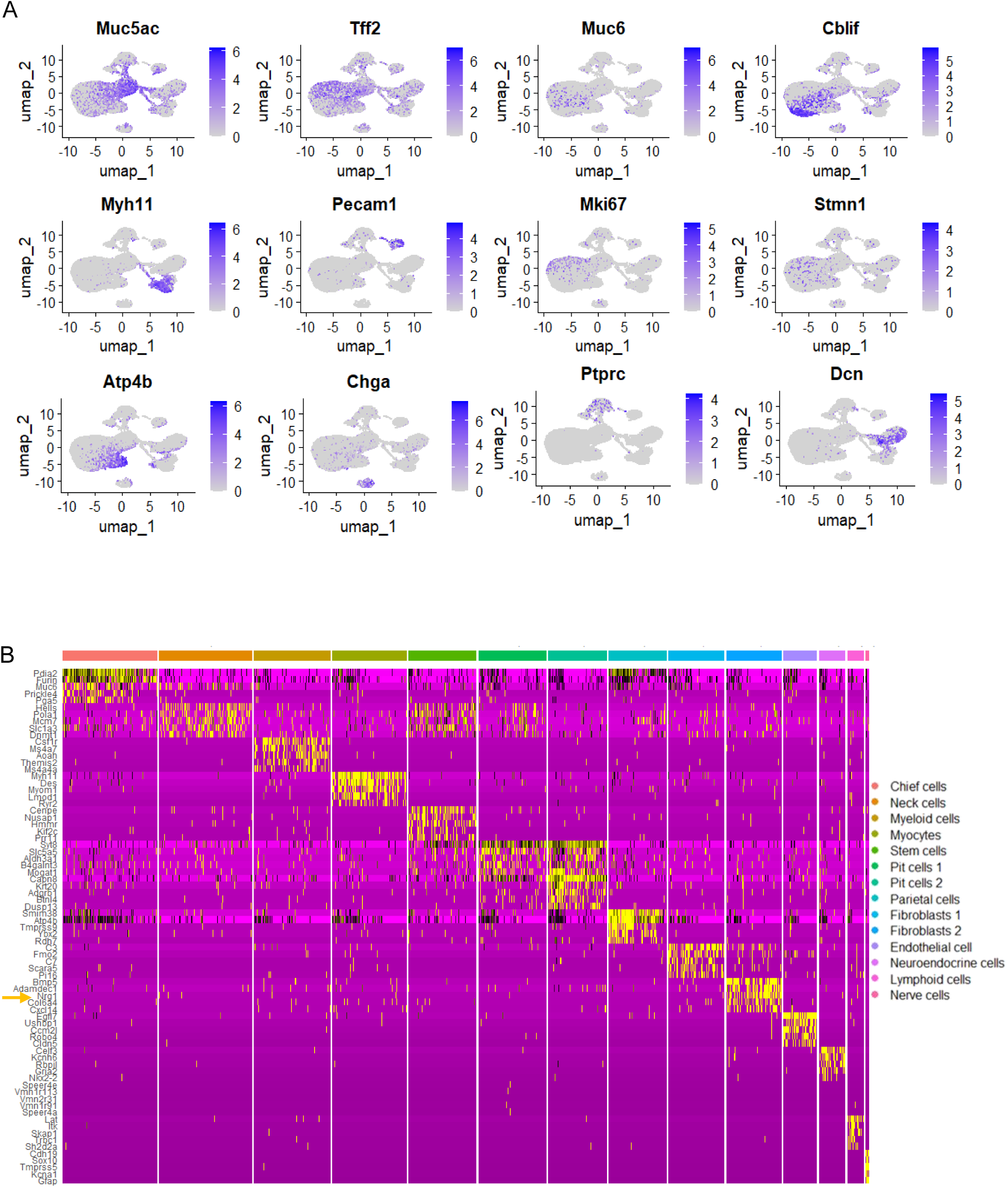
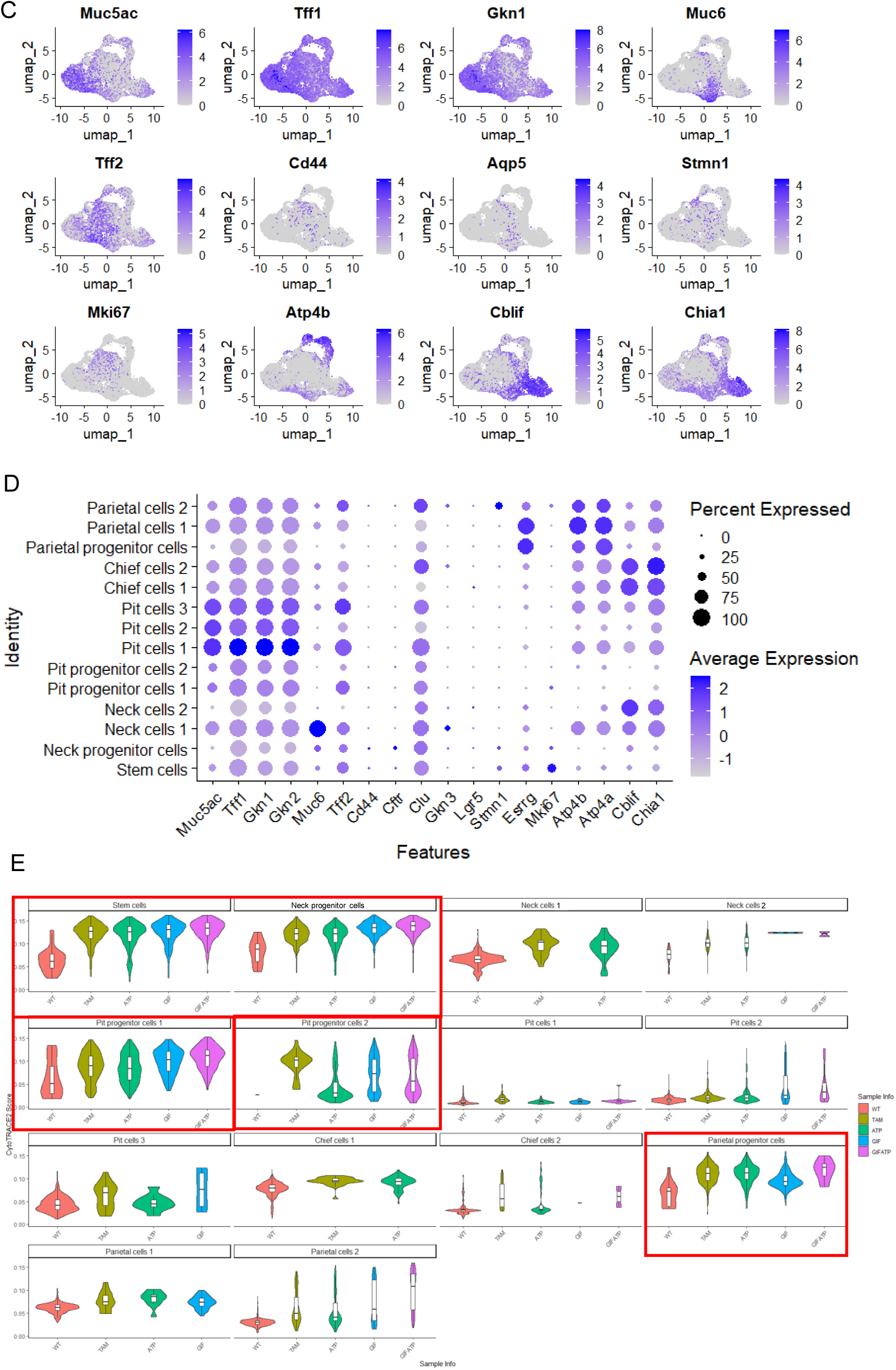
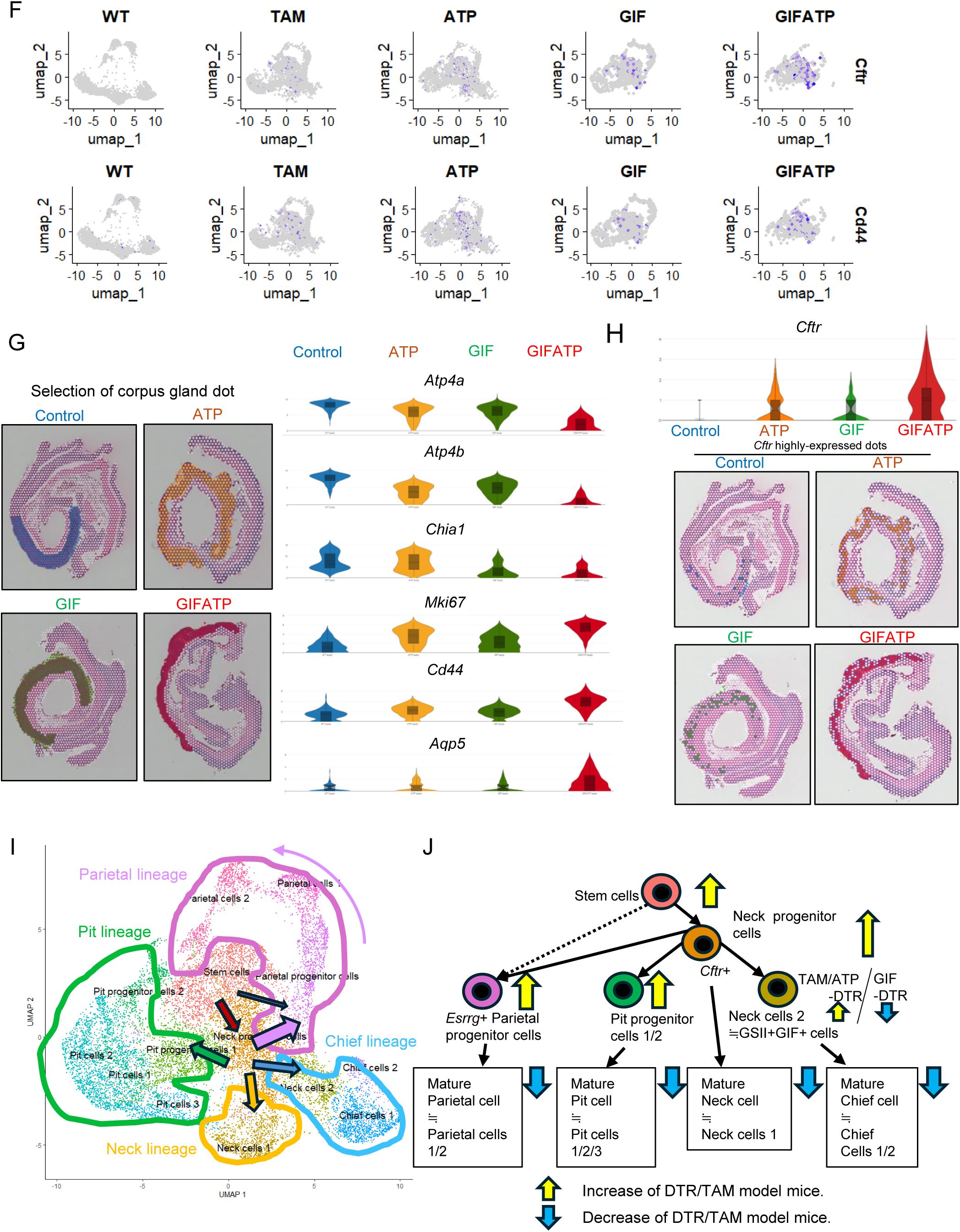
Single cell and spatial transcriptomic analyses of murine gastric injury models. (A) Feature plots of the indicated cell markers across the 14 clusters in the whole cohort. (B) Heat map of the 5 top DEGs across 14 clusters in the whole cohort. (C) Feature plots of the indicated cell markers across epithelial cell clusters. (D) Bubble plots showing the epithelial cell markers across epithelial cell clusters. (E) Violin plots showing the CytoTRACE2 scores across 14 epithelial clusters, split by WT, TAM, ATP, GIF, and GIFATP mice. (F) Feature plots showing the *Cftr* and *Cd44* expression, split by WT, TAM, ATP, GIF, and GIFATP mice. (G-H) The spatial transcriptomic Visium analysis with the dot of the corpus gland. (G) *Atp4b, ATP4b, Chia1, Mki67, Cd44*, and *Aqp5* expression across WT, ATP, GIF, and GIFATP mice shown as Violin plots. (H) *Cftr* expression across WT, ATP, GIF, and GIFATP mice shown as Violin plots and spatial gene distribution. (I-J) Schema of the gastric epithelial differentiation across WT, TAM, ATP, GIF, and GIFATP mice.

**Figure S3, related to Figure 3.**
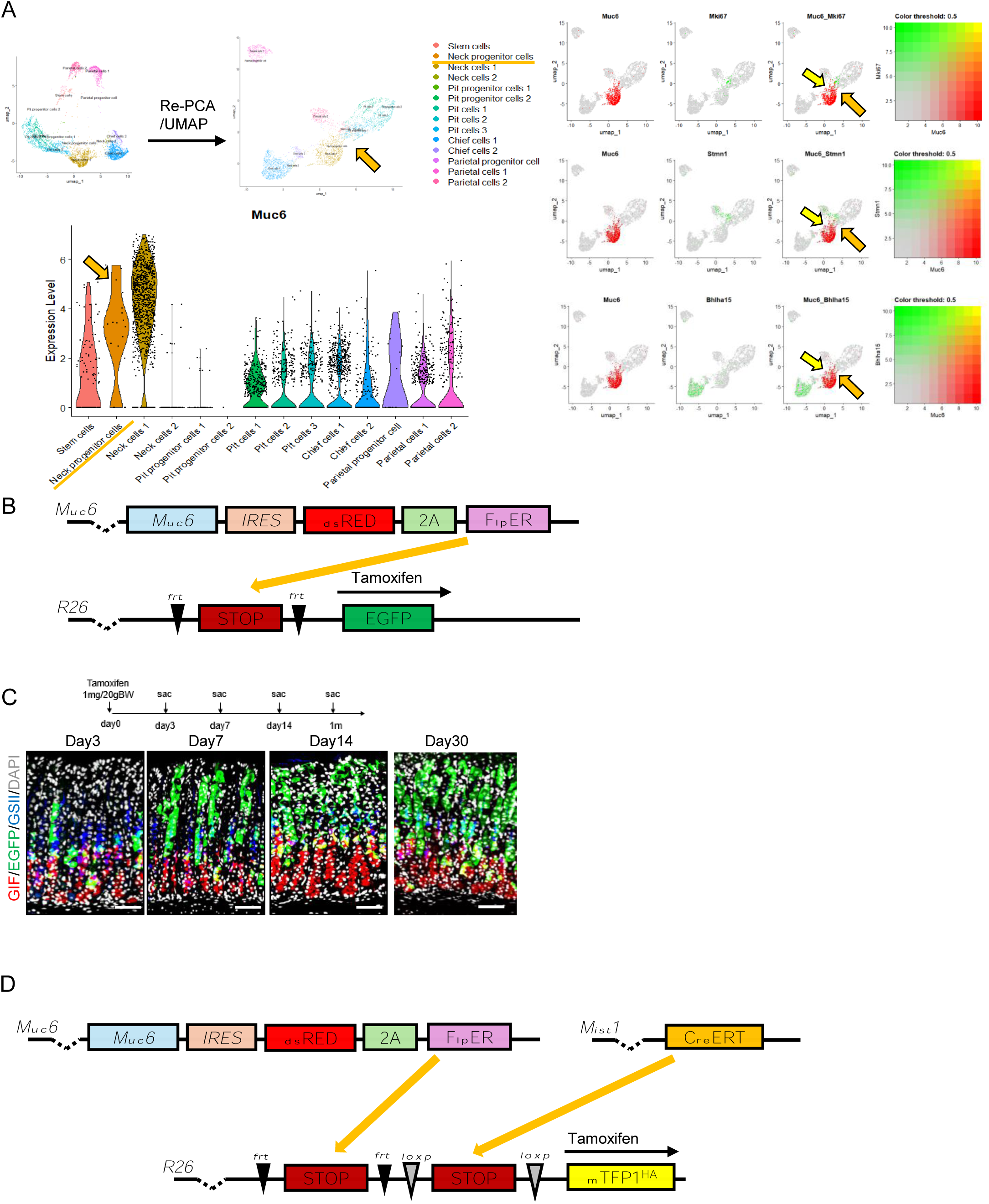
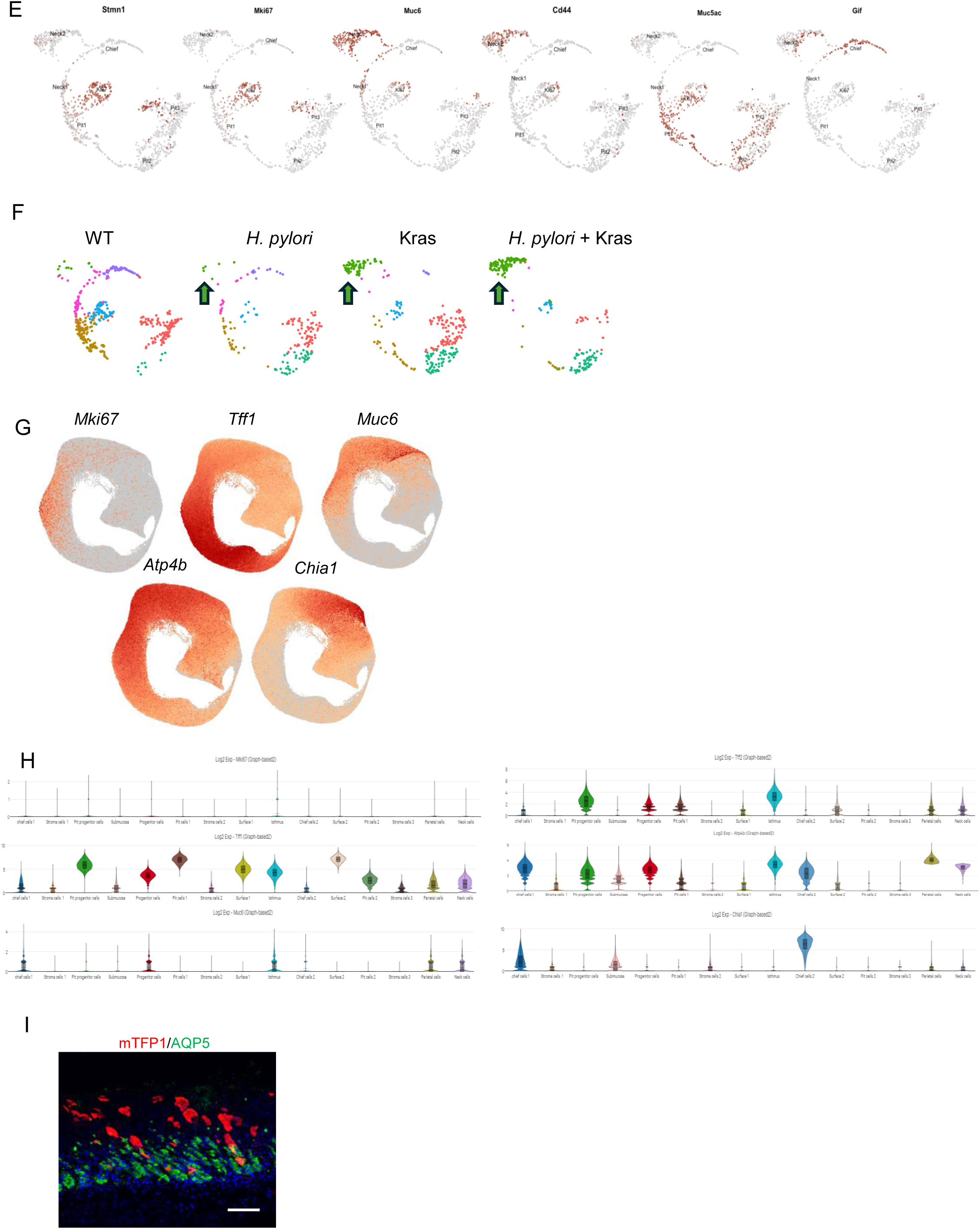
*Muc6*-expressing isthmus progenitors give rise to metaplasia in the stomach. (A) The epithelial cell clusters from WT mice were re-clustered and shown as a UMAP plot. *Muc6* expression was shown as violin plot across 14 epithelial cell clusters and co-expression analysis among *Muc6, Mki67, Stmn1*, and *Bhlha15* (Mist1) was performed. Orange and yellow arrows indicate neck progenitor cells and co-expressing cells respectively. (B) Gene constructs of *Muc6*-dsRED-FlpER; FSF-*EGFP* mice. (C) *Muc6*-dsRED-FlpER; FSF-*EGFP* treated with low-dose (50 mg/kg) tamoxifen. GIF (green)/EGFP (green)/GSII (blue) staining are shown. (D) Gene constructs of *Mist1*-CreERT; *Muc6*-dsRED-FlpER; LSL-FSF-mTfp1^HA^ mice. (E-F) scRNA-seq data with WT and *Mist1*-CreERT; LSL-*Kras*^G12D^ mice treated with or without *H. pylori.* (E) Feature plot showing the indicated cell markers. (F) UMAP plot of the gastric epithelial cell clusters split by the mouse groups. Green arrows show the SPEM clusters. (G-H) Spatial transcriptional Visium HD analysis with *Mist1*-CreERT and *Mist1*-CreERT; LSL-*Kras*^G12D^ mice 7 days after tamoxifen treatment. (G) Expression of cell markers shown in UMAP. (H) Expression of cell markers shown in Violin plots. (I) mTFP1(red)/AQP5(green) staining in *Mist1*-CreERT; *Muc6*-dsRED-FlpER; LSL-FSF-mTfp1^HA^ mice infected with *H. pylori* for 4 months. Scale bars; 100 μm. Mean ± S.E.M. *P < .05.

**Figure S4, related to Figure 4.**
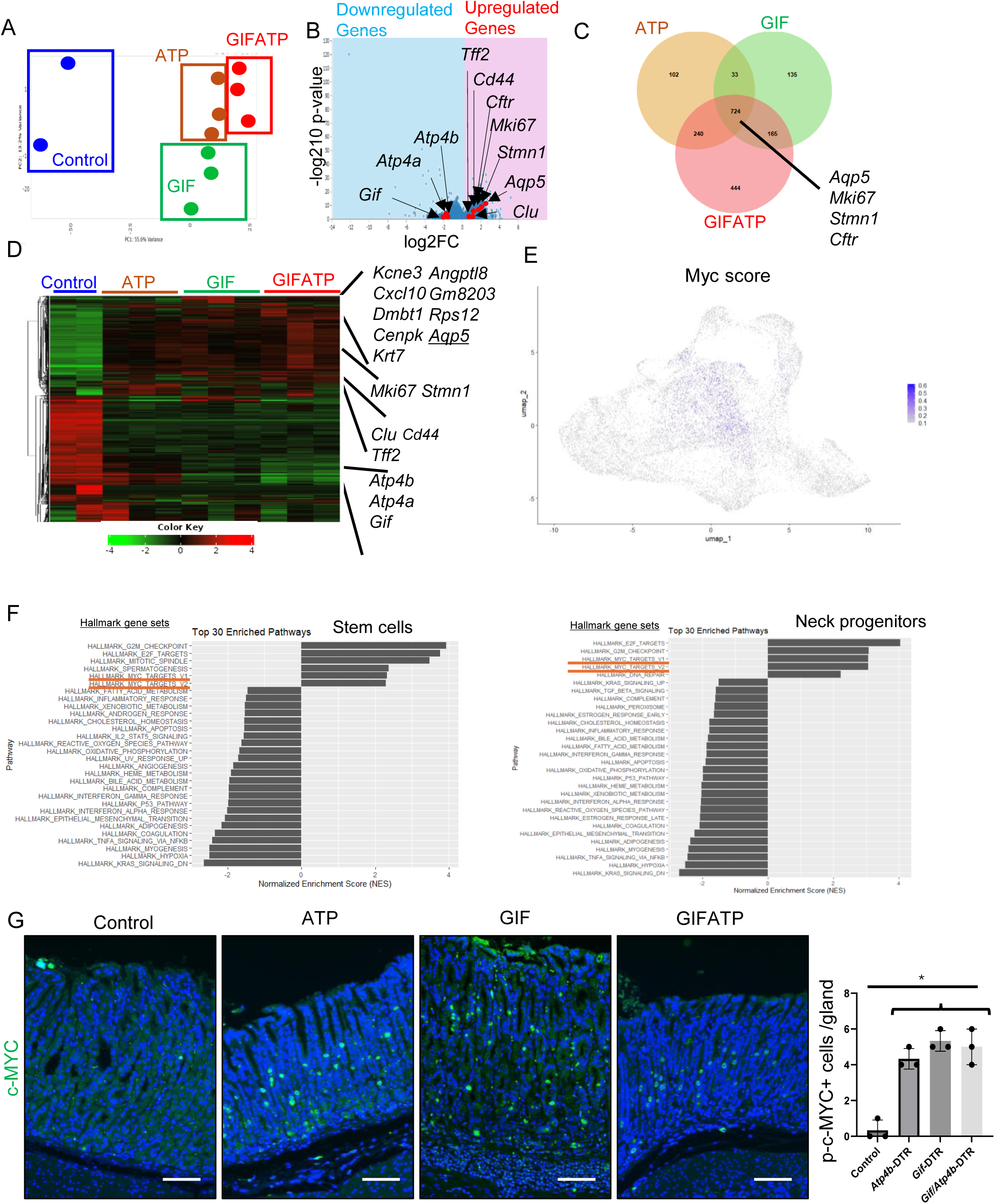
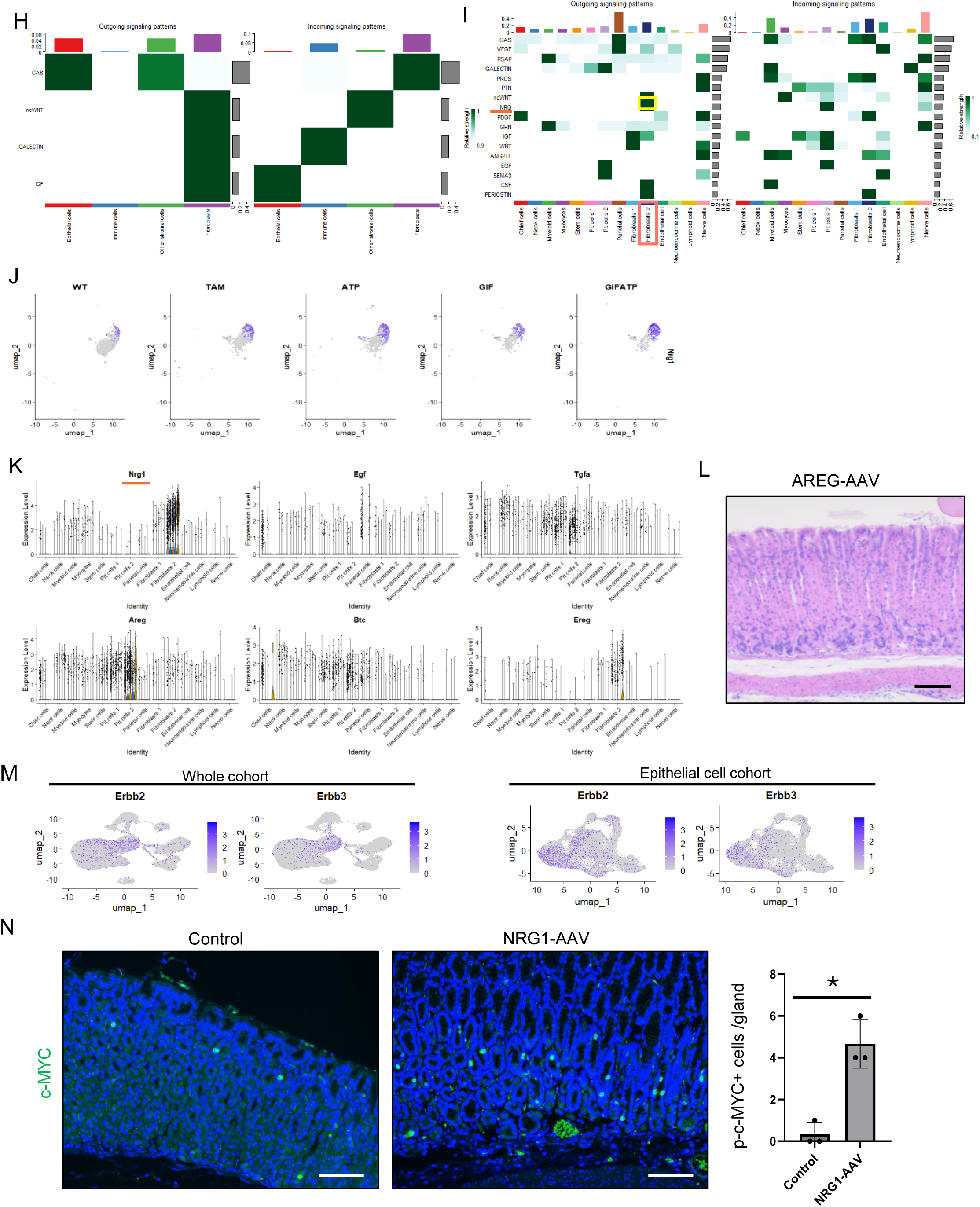
NRG1+ fibroblasts activate Myc pathway in isthmus progenitors following mucosal injury. (A-D) Bulk RNA sequence analysis. (A) PCA plot of control and DTR models. (B) Volcano plot showing *Clu, Mki67, Tff2, Cd44, Aqp5, Stmn1, Gif, Atp4a,* and *Atp4b* expression. (C) Venn diagram for upregulated DEGs in DTR models compared with control mice. (D) Heatmap of gene expressions. (E) Feature plot of gastric epithelial cell clusters in scRNAseq showing the Myc score. (F) fGSEA analysis of stem and neck progenitor cells with hallmark gene sets. (G) phospho-c-MYC staining and quantification in control, ATP, GIF and GIFATP mice (n=3 mice/group). (H) Heatmap of the outgoing and incoming interactions across epithelial, immune, fibroblast, and other stromal cells clusters. (I) Heatmap of the outgoing and incoming interactions across the whole clusters. (J) Feature plot showing the *Nrg1* expression in fibroblast clusters, split by WT, TAM, ATP, GIF, and GIFATP mice. (K) Violin plot showing the *Nrg1, Egf, Tgfa, Areg, Btc*, and *Ereg* expression across the whole clusters, split by WT, TAM, ATP, GIF, and GIFATP mice. (L) HE image of AREG-AAV treated mice (day 7). (M) Feature plot of *Erbb2* and *Erbb3* expression in total and gastric epithelial cell cohorts. (N) phospho-c-MYC staining and quantification in control- and NRG1-AAV treated mice (n=3 mice/group). Scale bars; 100 μm. Mean ± S.E.M. *P < .05.

**Figure S5, related to Figure 5.**
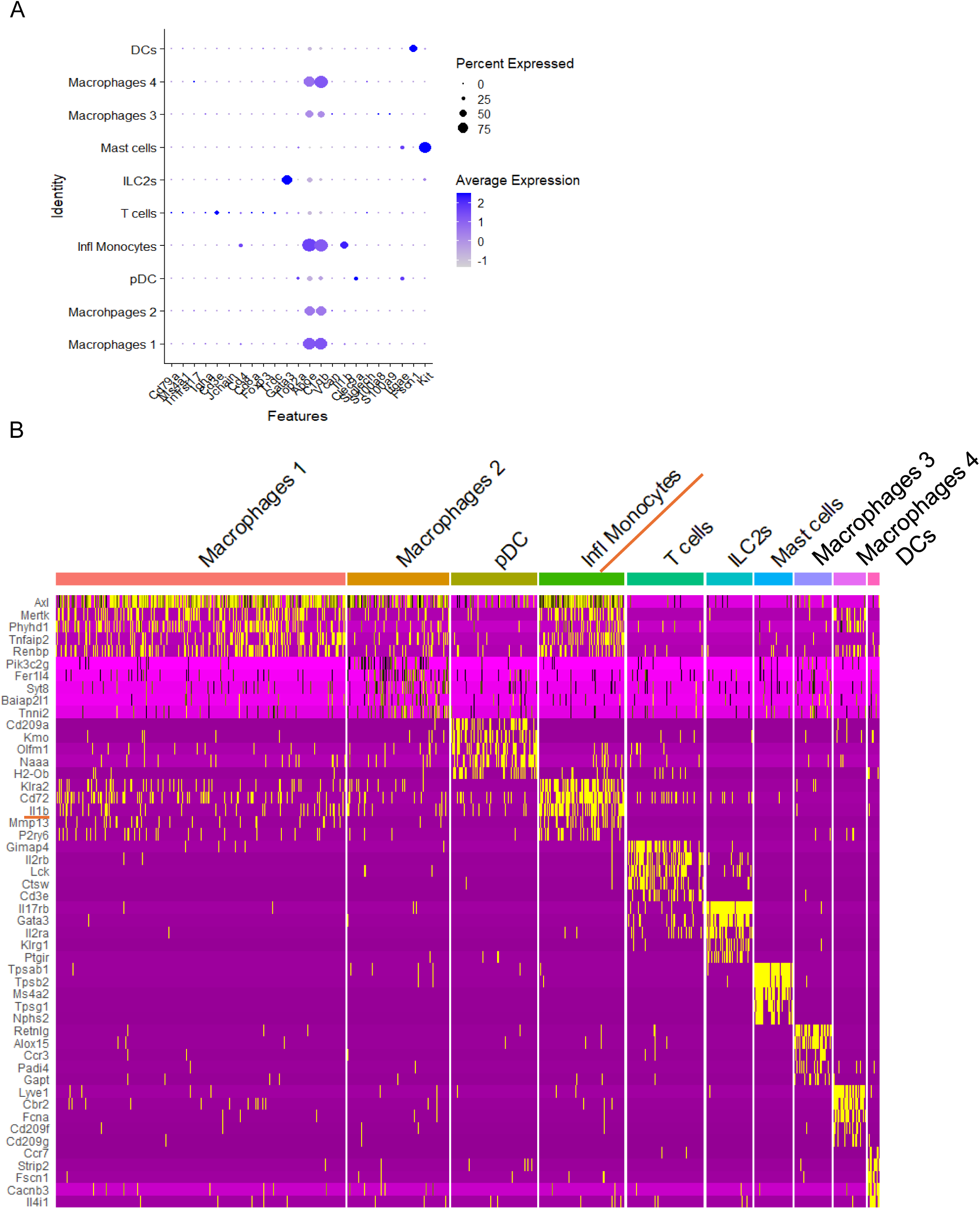
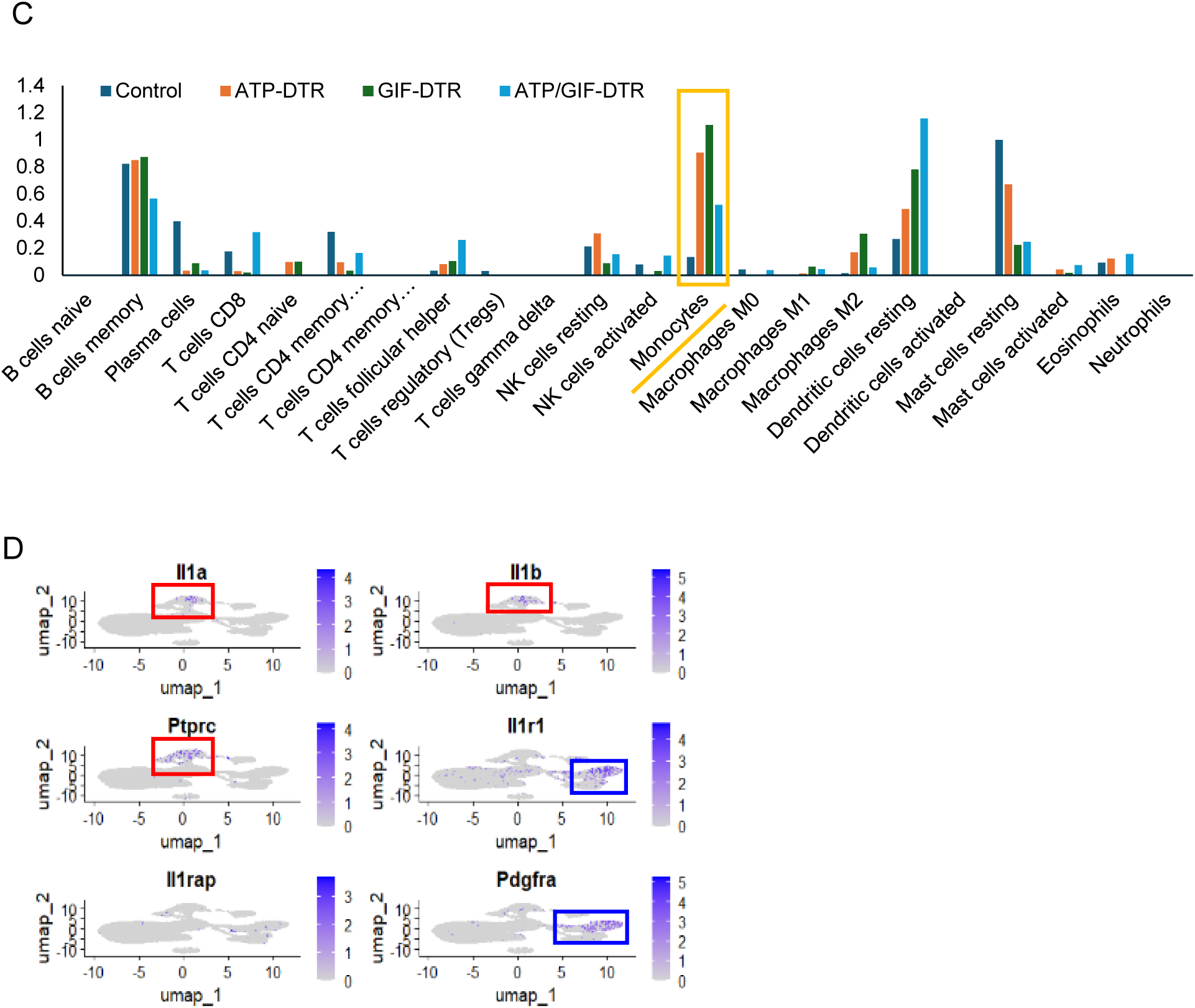
Inflammatory monocytes produce IL1 and interact with fibroblasts. (A) Dot plot of inflammatory cell markers across 10 inflammatory cell clusters. (B) Heatmap showing the top 5 DEGs across 10 inflammatory cell clusters. (C) Analysis of the inflammatory cells with CIBERSORTx in bulk RNAseq. (D) Feature plot of *Il1a*, *Il1b*, *Ptprc*, *Il1r1*, *Il1rap*, and *Pdgfra* expression in the whole single-cell RNA-seq samples.

**Figure S6, related to Figure 6.**
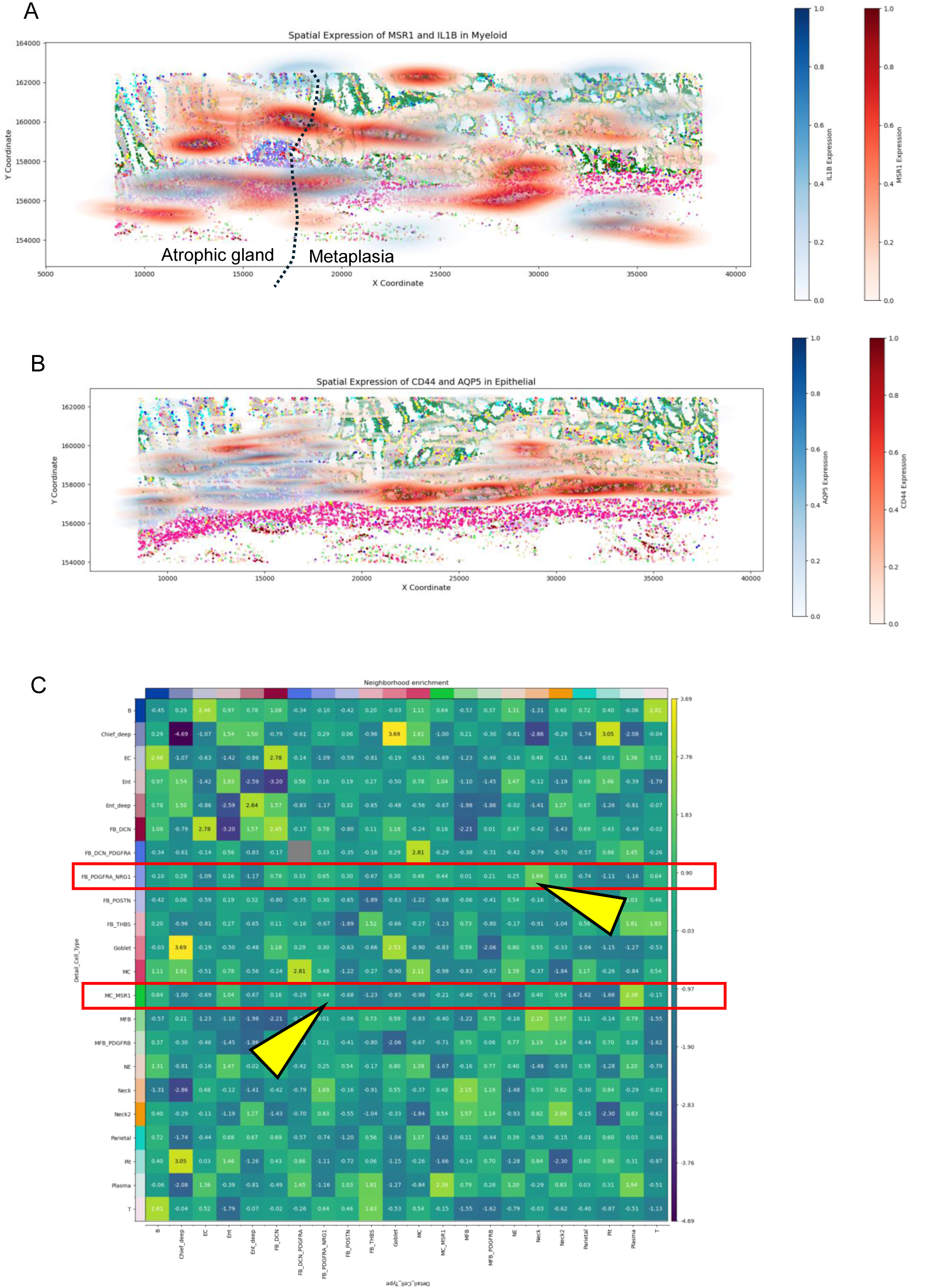
Spatial gene expression and cell-cell interaction analysis in human gastritis samples. (A-B) Kernel density plots with human spatial transcriptomic data. (A) *MSR1* and *IL1B* expressions derived from myeloid cells (B) *CD44* and *AQP5* expressions derived from epithelial cells. (C) Spatial proximity analysis restricted fundic glands adjacent to intestinal metaplasia. Yellow arrows show the proximity of NRG1+PDGFRA+ fibroblasts, neck cells, and MSR1+ monocytes. For this analysis, the Muscle cluster was excluded to focus on mucosal region. Additionally, similar cell clusters were merged: Parietal and Parietal2 into Parietal; Plasma and B_Plasma into Plasma; and Lowquality and Lymph_NS into Lowquality.

**Table S1.** Upregulated DEGs in DTR mice compared with control mice.

**Table S2.** Downregulated DEGs in DTR mice compared with control mice.

**Table S3.** Upregulated DEGs in the classified groups with Venn diagraph.

**Table S4.** GSEA results with HALLMARK gene sets in DTR mice compared with control mice.

## STAR★ Methods

### RESOURCE AVAILABILITY

#### Lead contact

Further information and requests for resources and reagents should be directed to and will be fulfilled by the lead contact, Yoku Hayakawa.

#### Materials availability

All the unique reagents generated in this study were available from the lead contact with a completed material transfer agreement.

#### Data and code availability

Transcriptomic data were deposited to GEO database (GSE292145). Further information and requests for resources and reagents should be directed to and will be fulfilled by the lead contact, Yoku Hayakawa.

### EXPERIMENTAL MODEL DETAILS

#### Human samples

Gastric tissue samples were collected from patients who underwent gastrectomy at The University of Tokyo Hospital. Formalin-fixed, paraffin-embedded (FFPE) tissue specimens were subsequently prepared and subjected to immunohistochemistry and spatial transcriptomic analyses.

This study was approved by the Research Ethics Committee of The University of Tokyo (approval number G3521), and written informed consent was obtained from all participating patients.

#### Mice

The *Loxp-STOP-Loxp (LSL)-Kras^G12D^*, *LSL*-TdTomato, *Mist1-creERT*, *Il1r1-/-*, and *Frt-STOP-Frt (FSF)*-EGFP mice were purchased from Jackson laboratories. The tetO-Cre^19^, *LSL-FSF*-mTFP1^HA^^41^ and *Msr1*-DTR mice^42^ were obtained from RIKEN animal resources. The *Tff1-*cre^43^, *Gpr30*-rtTA^19^, *Muc6*-dsRED-FlpER^44^ mice were described previously. The *Gif*-DTR, *Atp4b*-DTR, and *LSL-Cdx2* mice were generated as described in Method Details. All animal studies and procedures were approved by the ethics committees of the University of Tokyo and the Institute of Medical Science, Asahi Life Foundation were performed in compliance with institutional guidelines.

#### Organoid generation

Organoids were generated as described previously ^20,45^ . The mouse stomachs were cut longitudinally and washed with cold PBS, and the corpus parts were incubated in PBS containing 8 mM EDTA for 60 min on ice. Gland fractions were mechanically dissociated by cover slip, centrifuged at 900 rpm for 6 min at 4 °C, and diluted with advanced DMEM/F12 (Invitrogen, Waltham, MA, USA) containing B27, N2, 1 μM n-Acetylcysteine, 10 mM HEPES, penicillin/streptomycin, and Glutamax (all from Invitrogen). The glands were embedded in Matrigel and 500 glands/well were seeded in a pre-warmed 24-well plate. After Matrigel solidified, it was overlaid with advanced DMEM/F12 containing 50 ng/mL EGF (Invitrogen) and conditioned medium containing noggin, R-spondin, and Wnt3a. The medium was replaced twice weekly. Growth factors were removed when the proliferation was analyzed.

### METHOD DETAILS

#### Generation of Atp4b-DTR, Gif-DTR, and LSL-Cdx2 mice

We selected a sequence (5’-AGA GGA CAT GGC AGC CCT GC-3’) containing the start codon of the *Atp4b* as the sgRNA target. We inserted this sequence into pX330-mG plasmid, which carried both guide RNA and Cas9-mG expression units1. In the pBS-Atp4b-DTR-ZSgreen donor DNA, we placed the Chimeric intron2-HBEGF (human diphtheria toxin receptor gene, DTR)-2A-ZSgreen-bovine growth hormone polyadenylation (bGHpA) signal sequence between the 5’ and 3’ homology arms. The 5’-homology arm is the genomic region from 1,954 bp upstream to 1 bp upstream of the *Atp4b* start codon, and the 3’-homology arm is the genomic region from 1 bp downstream to 1,571 bp downstream of the *Atp4b* start codon. Above DNA vectors were isolated with FastGene Plasmid mini Kit (Nippon genetics, Tokyo, Japan) and filtrated by MILLEX-GV® 0.22 μm Filter unit (Merk Millipore, Darmstadt, Germany) for microinjection.The pregnant mare serum gonadotropin (5 units) and the human chorionic gonadotropin (5 units) were intraperitoneally injected into female C57BL/6J mice with a 48-h interval, and mated with male C57BL/6J mice. We collected zygotes from oviducts in mated female and mixture of pX330-mG (circular, 5 ng/μL, each) and pBS-Atp4b-DTR-ZSgreen (circular, 10 ng/μL) was microinjected into zygotes. Subsequently, injected zygotes were transferred into oviducts in pseudopregnant ICR female and newborns were obtained.

With the same strategy, we performed a knock-in into the *Gif* locus. A sequence (5’-AAG GGA GAC GTG GAC AGG CA-3’) containing the start codon of the *Gif* was selected as the sgRNA target. In the pBS-Gif-DTR donor DNA, we placed the Chimeric intron- DTR -rabbit globin polyadenylation (rGpA) signal sequence between the 5’ and 3’ homology arms. The 5’-homology arm is the genomic region from 1,804 bp upstream to 1 bp downstream of the *Gif* start codon, and the 3’-homology arm is the genomic region from 1 bp downstream to 1,708 bp downstream of the *Gif* start codon.

To generate LSL-*Cdx2* mice, CAG-loxP-STOP-LoxP-*Cdx2*-poly(A) cassettes was inserted into a pCALNL5 plasmid (DNA Bank, RIKEN Bio Resource Center, Ibaraki, Japan), followed by homologous recombination by injection into C57BL/6J blastocysts.

#### Treatment in vivo

To induce *H. pylori* infection in mice, 200 ul bacterial suspension containing 1 x 10^8^ CFU/ml *Helicobacter pylori (H. pylori;* PMSS1) was orally administered for three times in a week ^46–49^. For diphtheria toxin (DT) treatment, DT was administered via intraperitoneal injection at a dose of 50 µg/kg. Lineage-tracing experiments were performed as reported previously ^19^. For tamoxifen induction, 50-300 mg/kg of tamoxifen dissolved in 200 μl corn oil was orally administered as indicated in Figure legends. Doxycycline was administered in drinking water at a concentration of 0.2% for 3 days. For AAV injection experiments, 1.0 x 10^11^ genome copies of control, AREG-, and NRG1-expressing virus vectors were intravenously administered to mice, and mice were analyzed 7 days later. Mouse experiments were repeated at least two times, with at least 3 biological replicates per group.

#### Treatment in vitro

For organoid experiments, the recombinant NRG1 was added directly to the organoid medium (100 and 500 ng/ml). Organoid diameters were analyzed after 5 and 10 days. For fibroblast experiments, the recombinant IL1α and IL1β were added directly to the fibroblast medium (10 and 100 ng/ml). After 6 and 24 hours, fibroblasts were harvested.

#### Western blotting

Protein lysates were prepared from cultured cells, separated using sodium dodecyl sulfate-polyacrylamide gel electrophoresis, and transferred onto polyvinylidene difluoride membranes (Merck Millipore, Burlington, MA, USA). Membranes were probed with primary antibodies and incubated with secondary antibodies. Immunocomplexes were detected using an enhanced chemiluminescence system (Amersham Biosciences, Amersham, UK). The primary antibodies used were NRG1 (Abcam; 1:1000) and β-actin (Sigma Aldrich, St. Louis, MO, USA; 1:10000).

#### Histopathologic Analysis

Gastric tissues from mice were fixed in either 10% formalin or 4% paraformaldehyde, embedded in paraffin or optimum cutting temperature compound, respectively, and cut into 5-μm sections. Immunohistochemistry was performed as previously described ^50,51^. *In situ* hybridization was performed as described previously ^52^ or using an RNAscope kit (Advanced Cell Diagnostics, Newark, CA, USA) with *Cftr* and *Nrg1* probes. The average number of positive cells across 20 glands per mouse at the indicated sites was quantified.

#### Antibodies for Immunohistochemistry

The following primary antibodies were used: Ki67 (Cell Signaling Technology/Abcam/Biolegend; 1:200), H/K-ATPase (Santa Cruz Biotechnology, Santa Cruz; 1:200), pepsinogen I (BIO-RAD; 1:200), GIF (Columbia University, New York, NY, USA; 1:200), CD44v6 (Bio-RAD, Hercules; 1:200), CD44v9 (cosmo bio co. ltd; 1:200), GSII (Vector Laboratories, Newark, CA, USA; 1:200), TFF2 (Sino Biological, Beijing; 1:200), AQP5 (Sigma-Aldrich; 1:200), c-Myc (Abcam; 1:200), RFP (Rockland; 1:200), CDX2 (Abcam; 1:200), GFP (Abcam; 1:200), NRG1 (Abcam/Santa cruz; 1:200), Pdfgra (R and D; 1:200), HA (Roche; 1:200) and MSR1 (cell signaling Technology; 1:200).

##### Isolation of the stomach-specific fibroblasts

Mouse stomachs were excised and cut into approximately 1 cm segments. The tissues were rinsed in phosphate-buffered saline (PBS), then cut into ∼5 mm pieces and transferred into fresh Hank’s Balanced Salt Solution (HBSS) in 15 mL conical tubes. The tissues were vigorously washed with cold HBSS and the buffer was replaced with fresh HBSS for a total of eight washes.

Following the washes, the tissues were incubated in an enzymatic digestion solution containing 300 U/mL collagenase XI and 0.1 mg/mL dispase for 25 minutes at room temperature with gentle shaking. The digested tissues were centrifuged at low speed (4–10 × g) for 10 minutes. After centrifugation, the supernatant was discarded and the resulting pellet was transferred to 1.7 mL microcentrifuge tubes. The tissue clumps were thoroughly minced using scissors. Alternatively, tissue fragments were placed in a 6 cm dish and minced with a sterile blade.

The minced tissues were then washed in 15 mL tubes with 10 mL of DMEM containing 2% sorbitol (Sigma-Aldrich) by pipetting up and down. The suspension was centrifuged at 1500 rpm for 5 minutes, and the supernatant was discarded. This washing step was repeated five times. Subsequently, the cell suspensions were washed once more with 10 mL of DMEM and centrifuged at 10 × g for 2 minutes.

Finally, the resulting cell clumps were plated in 6-well plates. Tissue from 1 cm of stomach was sufficient to seed approximately six wells of a 6-well plate.

#### Bulk RNA sequence

DT was given to control and DTR model mice for 3 days (day 1, 2, and 3), then mice were sacrificed at day 4. RNA was extracted from the gastric mucosa of mice using a NucleoSpin RNA kit (MACHEREY-NAGEL GmbH & Co., Duren, Germany) or cultured using ISOGEN with a Spin Column kit (Nippon Gene, Tokyo, Japan). RNA was sequenced using the NovaSeq 6000 system. Data were analyzed on a log2 scale using the Subio platform (Kagoshima, Japan) or RaNA-seq website ^53^. The significance of the differential expression was analyzed using the Wald test (DESeq2). Differentially expressed transcripts were defined as those with a Wald test p-value <0.05, >2-fold upregulation, or < 0.5-fold downregulation. GSEA was performed as described previously^54^. The significance (P value) and false discovery rate (Q value) of the enrichment scores were determined using 1000 permutations of random gene sets of comparable sizes.

#### Spatial transcriptomic analysis (Visium)

DT was given to control and DTR model mice for 3 days (day 1, 2, and 3), then mice were sacrificed at day 4. The entire stomachs were removed, opened longitudinally, and fixed in 10 % formaldehyde in a swiss-roll structure for 24 hours. Fixed tissues were then embedded in paraffin blocks. After histological examination, one representative specimen from each group was selected and used for Visium analysis. RNA quality was assessed using the DV200 score, as recommended by 10x Genomics. Total RNA was extracted from FFPE tissue blocks using the RNeasy FFPE Kit (QIAGEN) following the manufacturer’s instructions. RNA quality was evaluated using the TapeStation system (Agilent). Only samples with a DV200 score >50% were used for subsequent spatial transcriptome analysis.

Mouse FFPE tissue blocks were sectioned at a thickness of 5 μm and mounted onto slides. The slides were processed according to the Visium CytAssist Spatial Gene Expression for FFPE protocol (10x Genomics, CG000520). Tissue sections were deparaffinized, stained with H&E, and imaged using the BZ-X810 microscope (KEYENCE). Following imaging, the sections were de-crosslinked to prepare for downstream analysis.

Library preparation was performed using the Visium Mouse Transcriptome Probe Kit (10x Genomics) following the manufacturer’s instructions, as outlined in the Visium CytAssist Spatial Gene Expression Reagent Kits User Guide (10x Genomics, CG000495). The optimal cycle number for index PCR was determined using the CFX Opus 96 Real-Time PCR System (Bio-Rad), and library quantification was performed using the TapeStation system (Agilent).

Sequencing was performed on the DNBSEQ-G400 platform (MGI) with a depth exceeding 50,000 read-pairs per spot, surpassing the manufacturer’s recommended sequencing depth. Raw sequencing reads were processed using SpaceRanger v.2.0.1 (10x Genomics) and aligned to the mm10-2020-A mouse reference genome.

#### Spatial transcriptomic analysis (Visium HD)

FFPE tissue sections were deparaffinized and stained with hematoxylin and eosin (H&E) according to the demonstrated protocol (CG000684, 10x Genomics). H&E imaging was obteined using BZ-X810 (Keyence). Probe hybridization was performed using Visium Mouse Transcriptome Probes v2. Visium HD spatial gene expression libraries were constructed according to the manufacturer’s protocol (CG000685, 10x Genomics). Sequencing was performed using the AVITI sequencer (Element Biosciences) with a 43/50 bp paired-end configuration. The obtained dataset was processed with Space Ranger v3.1.2 using the mouse reference mm10 v2020-A (10x Genomics).

#### Spatial transcriptomic analysis (CosMx 6K panel)

CosMx technology (Nanostring, Seattle, WA, USA), a high-plex spatial molecular imager1, was used to analyze three human gastric specimens: one fundic gland specimen and two metaplastic specimens that included both fundic and pyloric glands. B2M/CD298, PanCK, CD45, and CD3 antibodies and DAPI were used as morphological markers. The FFPE sections were processed as described by He et al ^55^. Cell segmentation was performed by combining image preprocessing and Cellpose^56^ neural network models, as detailed by He et al ^55^. Spatial transcriptome analysis was largely performed using python library scanpy ^57^ and squidpy^58^. For major cell type annotation, raw count data from the spatial transcriptome were normalized by total counts, scaling each cell’s total expression to a target sum of 10,000. After applying a log transformation using natural log and pseudocount 1, PCA was performed on the scaled data. Following dimensionality reduction, we computed the neighborhood graph using 15 nearest neighbors and 30 principal components with a cosine distance metric. We then generated a two-dimensional UMAP embedding and identified 20 clusters using the Louvain algorithm with default parameters in scanpy. Clusters were annotated as major cell types based on the expression of known marker genes (Plasma: IGHA1, IGKC, JCHAIN; Epithelial: KRT18, PIGR, PGA4, PSCA, TFF2, TFF1, LYZ, FABP1, MUC6; Lymphocyte: CD74, MHC I, HLA-DRB; Myeloid: CD74, HLA-DRB, C1QC, C1QB; Fibroblast: DES, TAGLN, COL1A1, POSTN, VIM; and Low-quality cells). We employed scVI^59^ to performed dimensionality reduction and batch effect removal to identify subclusters of major cell types with Sample_ID as batch key. After scVI processing, we performed Leiden algorithm with default parameters in scanpy and annotated subclusters by marker genes. Hallmark gene sets were retrieved from MSigDB ^60^ using msigdbr package. The gene symbols for each hallmark set were then compiled into a list structure for downstream analysis. Single-sample Gene Set Enrichment Analysis (ssGSEA) was performed on the normalized expression matrix using the gsva function from the GSVA package. To define spatial relationships among cells, we employed the sq.gr.spatial_neighbors function from the Squidpy package to capture local spatial connectivity between each cluster.

#### The analysis of single-cell RNA-seq

Droplet-based single-cell partitioning and single-cell RNA-Seq libraries were generated using the Chromium Fixed RNA Profiling Reagent Kits (10× Genomics) as per the manufacturer’s protocol based on the 10× GemCode proprietary technology.

Control and DTR model mice were given DT for 3 days (day 1, 2, and 3), then analyzed at day 4. For HDT samples, 300 mg/kg tamoxifen was given for every 3 days. Three fresh gastric corpus specimens from each group were pooled together and immediately frozen in liquid nitrogen. After fixing at least 25 mg of frozen tissue with 4% formaldehyde solution, cells were separated from the tissue using a gentleMACS Octo Dissociator (Miltenyi Biotec, Bergisch Gladbach, Germany) in a dissociation solution (Gibco RPMI RPMI 1640+ 0.2 mg/ml Liberase; Gibco, Waltham, MA, USA). The dissociated tissue was passed through a 30 µm filter to remove debris and undissociated tissue pieces.

The whole transcriptome probe pairs, consisting of the left-hand side (LHS) and right-hand side (RHS) of each targeted gene, were added to the fixed sample. The probe pairs hybridized to their complementary target RNA during overnight incubation.

After hybridization, the samples are washed three times, single-cell suspension at a density of some 3500 cells/µl was mixed with GEM master mix and immediately loaded together with Single Cell Fixed RNA Gel Bead and Partitioning Oil into a Chromium Next GEM Chip Q. The chip was then loaded onto a Chromium X (10× Genomics) for single-cell GEM generation and barcoding. Once the ligation and barcoding steps are completed, the GEMs are broken by the addition of Recovery Agent, and a PCR master mix is added directly to the post-GEM aqueous phase to pre-amplify the ligated products. The pre-amplified products are then cleaned up by SPRIselect (Backman Coulter).

The 10x barcoded, ligated probe products undergo indexing via Sample Index PCR. This, in turn, generates final library molecules that are cleaned up by SPRIselect, were incorporated into finished library which were compatible with Illumine next-generation short-read sequencing. The size profiles of the sequencing libray were examined by Agilent Bioanalyzer 2100 using a High Sensitivity DNA chip (Agilent).

The library were sequenced on Illumina NovaSeq X plus system using the NovaSeq X Series 10B Reagent Kit (Illumina) with a paired end, dual indexing (28/10/10/90-bp) format according to the recommendation by 10× Genomics.

Filtered count matrices were imported into R for further analysis using the Seurat package ^61^. We performed clustering according to the cell markers, depicted heatmap with top 5 DEGs, counted cell numbers in each cluster, checked the cell markers with violin plot, bubble plot, and feature plot, and conducted differential expression and gene set enrichment analysis on pseudobulk workflow by using “AggregateExpression” function in Seurat and R packages DESeq2 and fgsea.

For Figure 3G-I, scRNAseq data were obtained from previously published dataset^27^.

### QUANTIFICATION AND STATISTICAL ANALYSIS

Continuous variables were expressed as a means with 95% standard deviations, whereas categorical variables were expressed as numbers and frequencies (%). The differences between the means were compared using either Student’s t-test or the Wilcoxon test. Statistical significance was set at p < 0.05.

