## Supplementary material for "*Muc6*-expressing gastric isthmus progenitors contribute to regeneration and metaplasia supported by myeloid-mesenchymal interactions": Materials

**Key resources table**

| REAGENT or RESOURCE | SOURCE | IDENTIFIER |
| --- | --- | --- |
| Antibodies | | |
| Rabbit monoclonal anti-β-actin (HRP conjugated) | cell signaling technology | 5125S |
| Mouse monoclonal anti-H/K-ATPase | Santa cruz biotechnology | Sc-374094 |
| Rabbit monoclonal anti-GIF | Columbia university | NA |
| Rabbit monoclonal anti-Ki67 | cell signaling technology | 12202S |
| Rabbit monoclonal anti-Ki67 | Abcam | ab16667 |
| Rat monoclonal anti-Ki67 | Biolegend | 151202 |
| Rabbit polyclonal anti-TFF2 | sino biological, | 200208-T08 |
| Goat monoclonal anti-CD44v6 | Bio RAD | MCA1967 |
| Rabbit polyclonal anti-RFP | Rockland | 600-401-379 |
| Goat polyclonal anti-RFP | Rockland | 200-101-379 |
| Goat polyclonal anti-GFP | Abcam | Ab5450 |
| Mouse monoclonal anti-PG1 | BIO-RAD | 7240-1009 |
| Rat monoclonal anti- CD44v9 | cosmo bio co. ltd | LKG-M003 |
| FITC-conjugated Griffonia（Bandeiraea）simplicifolia Lectin 2 | VECTOR LABORATORIES | FL-1211 |
| Rabbit monoclonal anti-MSR1 | cell signaling technology | 91119T |
| Rabbit polyclonal anti-NRG1 | Abcam | ab191139 |
| Mouse monoclonal anti-NRG1 | Santa Cruz | sc-393006 |
| Goat polyclonal anti-PDGFRA | R and D | AF1062 |
| Rabbit monoclonal anti-AQP5 | Sigma-Aldrich | HPA065008 |
| Rabbit monoclonal c-Myc | Abcam | ab185656 |
| Rabbit monoclonal CDX2 | Abcam | ab76541 |
| Rat monoclonal HA | Roche | 11867423001 |
| Bacterial and virus strains | | |
| pAAV[Exp]-EF1A>mNrg1 | Vectorbuilder | VB241211-1774ryy |
| pAAV[Exp]-EF1A>mAreg | Vectorbuilder | VB250130-1671yus |
| Chemicals, peptides, and recombinant proteins | | |
| Recombinant Mouse IL-1α | Bio legend | 575006 |
| Recombinant Mouse IL-1β | Bio legend | 575106 |
| Recombinant NRG1/HRG1 Protein, CF | R and D | 5898-NR-050 |
| Critical commercial assays | | |
| RNAscope(R) 2.5 HD Reagent Kit- RED | ACD | ADC-322350 |
| NucleoSpin RNA | MACHEREY-NAGEL | 740955.10 |
| ISOGEN with a Spin Column kit | Nippon Gene | 318-07511 |
| VECTASTAIN Elite ABC Kit | VECTOR LABORATORIES | PK-6100 |
| Deposited data |  |  |
| Raw and analyzed data | This paper | GSE292145 |
| Experimental models: Organisms/strains | | |
| Mouse: B6.129-Bhlha15tm3(cre/ERT2)Skz/J | The Jackson Laboratory | IMSR_JAX:029228 |
| B6.129S4-Krastm4Tyj/J | The Jackson Laboratory | IMSR_JAX:008179 |
| B6.Cg-Gt(ROSA)26Sortm14(CAG-tdTomato)Hze/J | The Jackson Laboratory | IMSR_JAX:007914 |
| B6.129S7-Il1r1tm1Imx/J | The Jackson Laboratory | IMSR_JAX:003245 |
| STOCK Gt(ROSA)26Sortm1.2(CAG-EGFP)Fsh/Mmjax | The Jackson Laboratory | MMRRC_032038-JAX |
| B6;129S6-Gt(ROSA)26Sor<tm1(CAG-mTFP1)Imayo> | Riken BioResource Research Center | RBRC05146 |
| C57BL/6-Msr1<tm1(HBEGF)Mtka> | Riken BioResource Research Center | RBRC06357 |
| B6.Cg-Tg(tetO-cre)1Tfur | Riken BioResource Research Center | RBRC05563 |
| Tff1-Cre | Original generation at Tokyo university | PMID: 30168144 |
| Muc6-dsRED-FlpER | Original generation at Tokyo university | PMID: 38583723 |
| Gpr30-rtTA | Original generation at Tokyo university | PMID: 32032583 |
| Gif-DTR | Original generation at Tokyo university | This paper |
| Atp4b-DTR | Original generation at Tokyo university | This paper |
| LSL-Cdx2 | Original generation at Tokyo university | This paper |
| Software and algorithms | | |
| GraphPad Prism version 8.0.0 for Windows | GraphPad | https://www.graphpad.com/guides/prism/latest/user-guide/citing_graphpad_prism.htm |
| CLC Genomics Workbench software v7.5 | Qiagen | https://digitalinsights.qiagen.com/products-overview/discovery-insights-portfolio/analysis-and-visualization/qiagen-clc-genomics-workbench/ |
